# Neuronal adaptation and optimal coding in economic decisions

**DOI:** 10.1101/147900

**Authors:** Aldo Rustichini, Katherine E Conen, Xinying Cai, Camillo Padoa-Schioppa

**Author notes:** Present address: NYU Shanghai, Shanghai, China. Correspondence: Camillo Padoa-Schioppa, Ph.D., Department of Neuroscience, Washington University in St Louis, Campus Box 8108.

## Abstract

During economic decisions, neurons in orbitofrontal cortex (OFC) encode the values of offered goods. Importantly, their responses adapt to the range of values available in any given context. Prima facie, range adaptation seems to provide an efficient representation. However, uncorrected adaptation in the encoding of offer values would induce arbitrary choice biases. Thus a fundamental and open question is whether range adaptation is behaviorally advantageous. Here we present a theory of optimal coding for economic decisions. In a nutshell, the representation of offer values is optimal if it ensures maximal expected payoff. In this framework, we examine the activity of *offer value* cells in non-human primates. We show that their firing rates are quasi-linear functions of the offered values, even when optimal tuning functions would be highly non-linear. Most importantly, we demonstrate that for linear tuning functions range adaptation maximizes the expected payoff, even if the effects of adaptation are corrected to avoid choice biases. Thus value coding in OFC is functionally rigid (linear tuning) but parametrically plastic (range adaptation with optimal gain). Importantly, the benefit of range adaptation outweighs the cost of functional rigidity. While generally suboptimal, linear tuning may facilitate transitive choices.

## Introduction

The relation between optimal coding and neuronal adaptation has often been examined in sensory systems^1-3^. One emerging concept is that some aspects of sensory coding are plastic and optimally adapting to the current behavioral context^4-10^ while other aspects are optimized for the distribution of natural stimuli and thus presumably rigid^4,11,12^. Importantly, uncorrected adaptation makes firing rates intrinsically ambiguous (coding catastrophe)^13,14^. Thus neuronal adaptation at any processing stage must be corrected at later stages^15^ or ultimately results in impaired behavioral performance^16,17^. Here we examined the relation between optimal coding and neuronal adaptation in the neural circuit underlying economic decisions. This circuit presents interesting analogies and notable differences with sensory systems.

Choosing between two goods entails computing and comparing their subjective values. Evidence from lesions and neurophysiology indicates that these mental operations engage the OFC^18-20^. Experiments in which rhesus monkeys chose between different juices identified three groups of neurons in this area. *Offer value* cells encode the values of individual goods and are thought to provide the primary input to the decision. Conversely, *chosen juice* cells and *chosen value* cells represent the binary choice outcome and the value of the chosen good^21,22^. Here we focus on *offer value* cells. Because they constitute the input layer of the decision circuit, these neurons are in some ways analogous to sensory cells. However, the behavioral goal subserved by *offer value* cells differs from that subserved by neurons in sensory systems. For the purpose of accurate perception, sensory neurons are optimally tuned if they transmit maximal information about the external world^1-3^. This goal is achieved if tuning functions match the cumulative distribution of the encoded stimuli^2,7^. In contrast, the purpose of a subject performing economic decisions is to maximize the payoff (i.e., the chosen value). Thus *offer value* cells are optimally tuned if they ensure maximal expected payoff.

As in sensory neurons, the issue of optimal tuning in *offer value* cells is intertwined with that of neuronal adaptation. Previous work indicated that *offer value* cells have linear responses and undergo range adaptation^21,23^. In other words, their tuning slope is inversely proportional to the range of values offered in any behavioral context^23-25^. Prima facie, a range adapting representation seems computationally efficient. However, it was shown that uncorrected adaptation in *offer value* cells would result in arbitrary choice biases^26^ conceptually analogous to the coding catastrophe discussed for sensory systems^14^. Thus it remains unclear whether range adaptation is behaviorally advantageous.

Here we present a series of theoretical and experimental results shedding new light on the nature of value coding and the role played by neuronal adaptation in economic decisions. Behavioral and neuronal data were collected in two experiments in which monkeys chose between different juices offered in variable amounts. First, we show that *offer value* tuning functions are quasi-linear and do not match the cumulative distributions of offered values. Second, we show that range adaptation is corrected within the decision circuit to avoid choice biases. Third, we develop a theory of optimal coding for economic decisions. We thus demonstrate that range adaptation, corrected to avoid choice biases, ensures maximal expected payoff. Fourth, confirming theoretical predictions, we show that payoff and value range are inversely related in the experiments. Fifth, we demonstrate that linear response functions were in fact suboptimal in our experiments. Hence, linearity is a rigid property of value coding not subject to contextual adaptation. Finally, we show that the benefit afforded by range adaptation outweigh the cost imposed by functional rigidity (linear tuning).

## Results

### Relative value, choice variability and expected payoff

In Exp.1, monkeys chose between two juices (A and B, with A preferred) offered in variable amounts (Fig.1ab). The range of quantities offered for each juice remained fixed within a session, while the quantity offered on any given trial varied pseudo-randomly. Monkeys’ choices generally presented a quality-quantity trade-off. If the two juices were offered in equal amounts, the animal would generally choose A (by definition). However, if sufficiently large quantities of juice B were offered against one drop of juice A, the animal would choose B. The "choice pattern" was defined as the percentage of trials in which the animal chose juice B as a function of the offer type. In each session, the choice pattern was fitted with a sigmoid function, and the flex of the sigmoid provided a measure for the relative value of the two juices, referred to as *ρ* (see Methods). The relative value allows to express quantities of the two juices on a common value scale. In one representative session, we measured *ρ* = 4.1 (Fig.1b).

**Figure 1.**
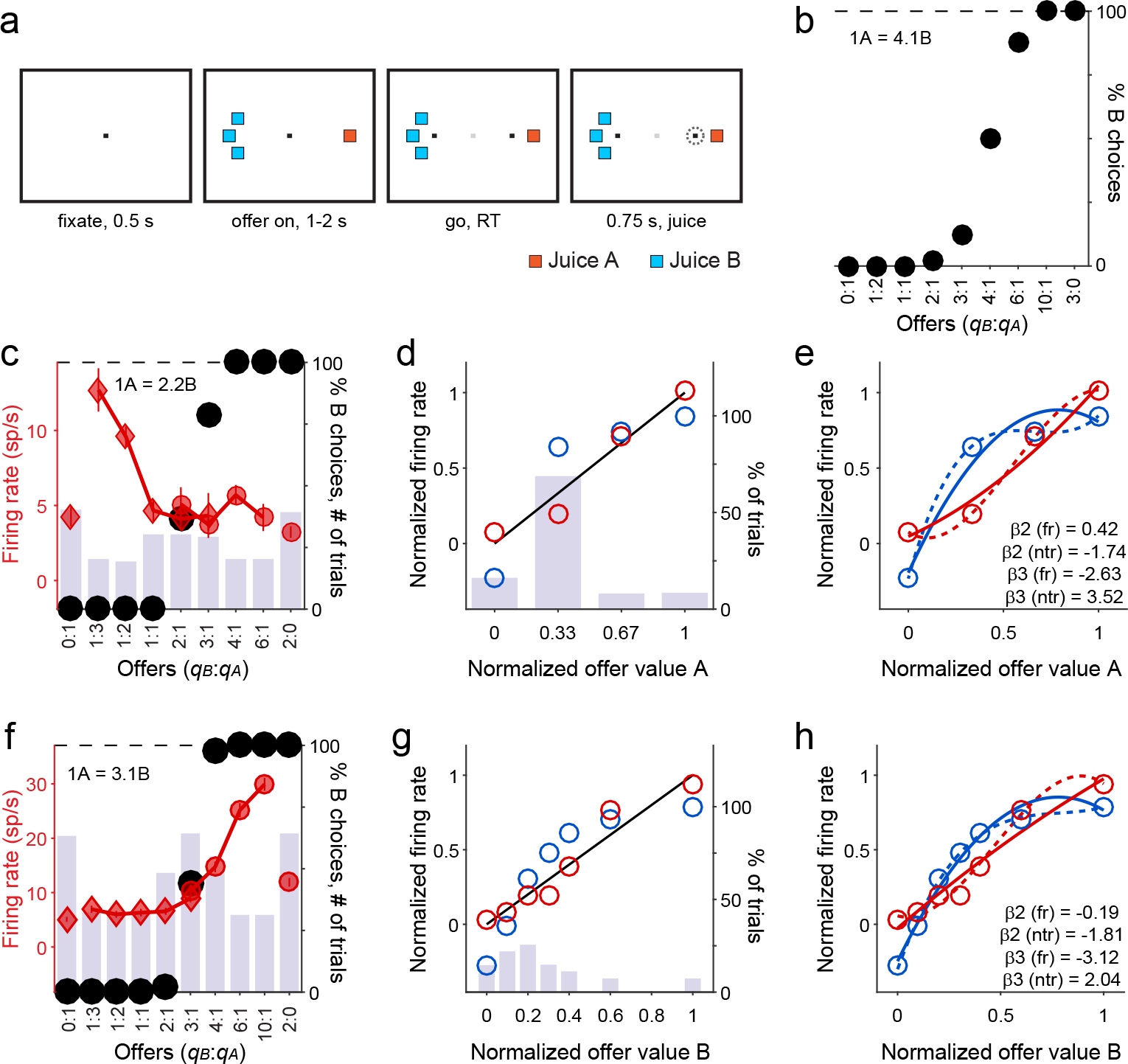
Quasi-linear coding of offer values, individual responses. **a.** Task design (see Methods). **b.** Example of choice pattern. The x-axis represents different offer types, ranked by the ratio *q_B_*:*q_A_*. Black dots represent the percent of "choice B" trials. **c.** Example *offer value A* response. Black dots represent the choice pattern. The histogram illustrates the number of trials presented for each offer type. Red symbols represent firing rates ± SEM (diamonds and squares for "choice A" and "choice B", respectively). The y-axis on the left refers to firing rates. The y-axis on the right refers both to the number of trials (histogram) and to the choice pattern (black symbols). **d.** Comparing firing rates and ntrials_CDF_. Same response as in (c). The x-axis represents normalized quantity levels of juice A (Methods). The histogram illustrates the percent of trials for each quantity level. This session included 247 trials, and juice A was offered at quantity levels 0 (39 trials, 16%), 1 (169 trials, 68%), 2 (19 trials, 8%), and 3 (20 trials, 8%). Note that low quantity levels were over-represented. Blue circles represent the cumulative distribution function (ntrials_CDF_). The y-axis on the right refers both to the number of trials (histogram) and to ntrials_CDF_ (blue circles). Red circles represent firing rates. Here each neuronal data point is an average across all the trials with given quantity level (not across a single trial type). The y-axis on the left refers to normalized firing rates. Limits on the y-axes were set such that the same line (black) represents the linear fit for firing rates and for ntrials_CDF_. (Since all measures are normalized, this is the identity line.) **e.** Curvature of firing rates and ntrials_CDF_. Same data points as in (d). Continuous and dotted lines are the result of the quadratic and cubic fit, respectively. **f-h.** Example *offer value B* response.

Choice patterns often presented some variability. For example, consider in Fig.1b offers 6B:1A. In most trials, the animal chose juice B, consistent with the fact that the value of 6B was higher than the value of 1A. However, in some trials, the animal chose the option with the lower value. Similarly, in some trials, the animal chose 3B over 1A. Intuitively, choice variability is high when the sigmoid is shallow. Thus in each session, the steepness of the fitted sigmoid, referred to as *η*, quantified the (inverse of) choice variability (see Methods).

In any given trial, we define the payoff as the value chosen by the animal. Thus given a set of offers and a sigmoid function, the expected payoff is equal to the chosen value averaged across trials. Importantly, the expected payoff is inversely related to choice variability, and thus directly related to the steepness of the sigmoid. When the sigmoid is steeper, choice variability is lower, and the expected payoff is higher; when the sigmoid is shallower, choice variability is higher, and the expected payoff is lower. Notably, the relative value of two juices is entirely subjective. In contrast, a key aspect of the expected payoff is objective: Given a set of offers, a relative value and two sigmoid functions, the steeper sigmoid yields higher expected payoff.

### Quasi-linear coding of offer values

While animals performed the task, we recorded the activity of individual neurons in the central OFC. Firing rates were analyzed in multiple time windows. In each session, an "offer type" was defined by a pair of offers (e.g., [1A:3B]); a "trial type" was defined by an offer type and a choice (e.g., [1A:3B, B]); a "neuronal response" was defined as the activity of one cell in one time window as a function of the trial type. Earlier work showed that different responses encoded variables *offer value*, *chosen value* and *chosen juice*^21,22^. Unless otherwise indicated, the present analyses focus on *offer value* responses.

Previous studies failed to emphasize that the tuning of *offer value* cells was quasi-linear even though the distribution of values was highly non-uniform. To illustrate this point, we identified for each *offer value* response the quantity levels for the corresponding juice, and we calculated the number of trials in which each quantity level had been presented to the animal within the session. For example, Fig.1c illustrates one *offer value A* response. In this session, juice A was offered in quantity levels (number of trials): 0 (39), 1 (169), 2 (19) and 3 (20). For each set of trials, we computed the mean firing rate of the cell. In addition, we computed the cumulative distribution function for the number of trials (ntrials_CDF_) as a function of the quantity level. By analogy with sensory systems^2^, neurons encoding ntrials_CDF_ would provide maximal information about the offer values. Notably, firing rates and ntrials_CDF_ were highly correlated. For both of them, a linear regression on the quantity level provided a reasonably good fit (Fig.1d). However, the non-uniform distribution of trials induced a curvature in ntrials_CDF_. Similarly, each neuronal response taken alone always presented some curvature. To assess whether and how the curvature in neuronal responses was related to the curvature in ntrials_CDF_, we normalized both firing rates and ntrials_CDF_ (Methods). We thus fit each set of data points with a 2D polynomial, which provided a coefficient for the quadratic term (β_2_). Separately, we fit each set of data points with a 3D polynomial, which provided a coefficient for the cubic term (β_3_; Fig.1e; see also Fig.1f-h).

Because trials at lower quantity levels were over-represented in the experiments, we generally measured β2, ntrialsCDF<0 and β3, ntrialsCDF>0. In contrast, β2, firing rate and β3, firing rate varied broadly across the population, and their distributions were fairly symmetric around zero (Fig.2). In other words, neuronal response functions were, on average, quasi-linear. These results held true for individual monkeys, in each time window, and independently of the sign of the encoding (Fig.S1). Similar results were also obtained for *chosen value* responses (Fig.S2).

**Figure 2.**
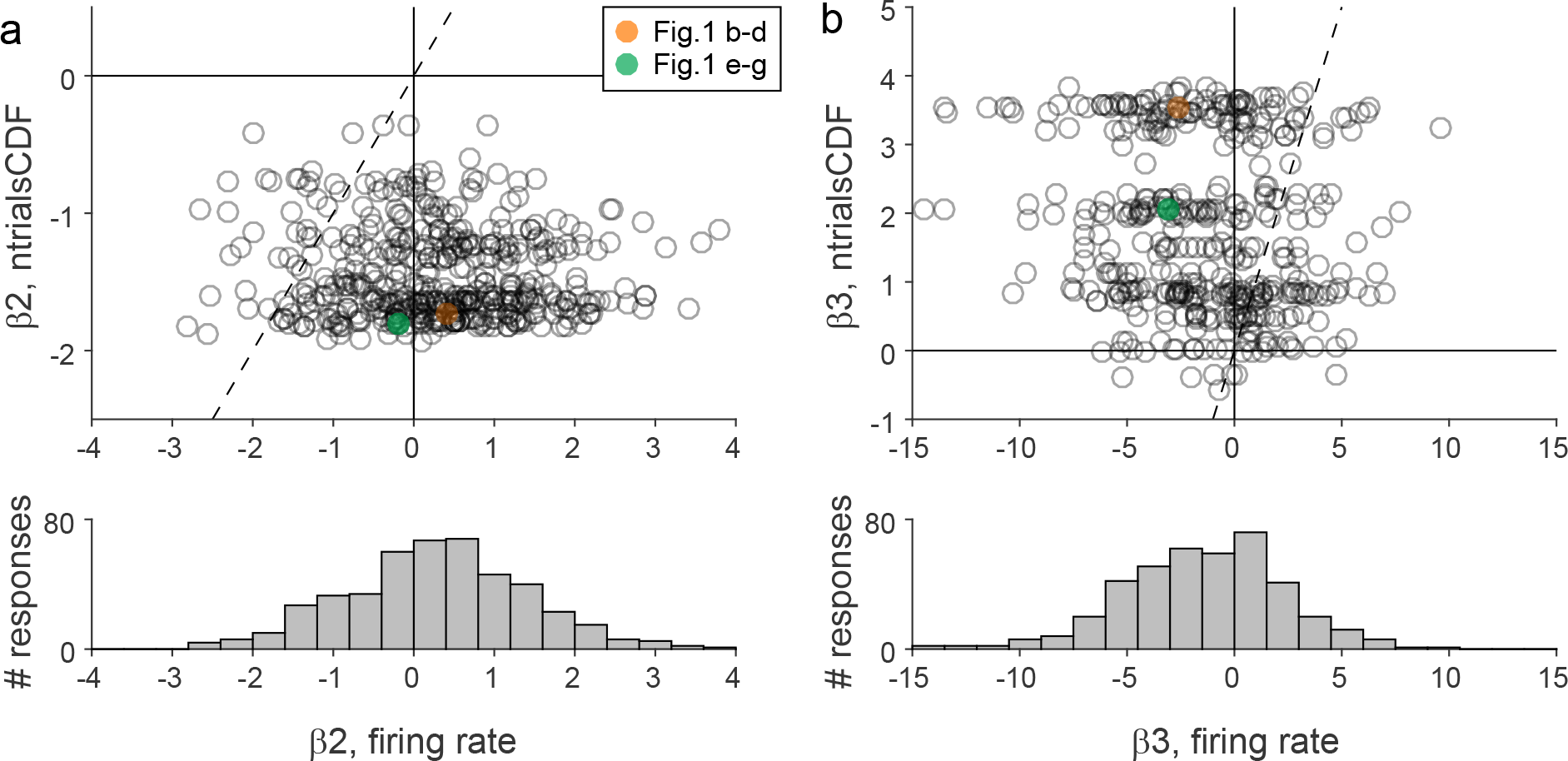
Quasi-linear coding of offer values, population analysis (N = 447). **a.** Quadratic term. Each data point in the scatter plot represents one response. The x-axis and y-axis represent β_2, firing rate_ and β_2, ntrials CDF_, respectively. The dotted line represents the identity line, and the responses illustrated in (b-g) are highlighted. Since low values were always over-represented, generally β_2, ntrials CDF_<0. In contrast, measures for β_2, firing rate_ were broadly scattered above and below zero (see histogram). **b.** Cubic term. Same conventions as in (a). Generally, β_3, ntrials CDF_>0. In contrast, measures for β_3, firing rate_ were broadly scattered above and below zero. Note that on average across the population, measures for β_2, firing rate_ were close to, but significantly above zero (mean(β_2, firing rate_) = 0.28; p<10^-6^, t-test). Conversely, measures for β_3, firing rate_ were close to, but significantly below zero (mean(β_3, firing rate_) = -1.42; p<10^-12^, t-test). We return to this point in the last section of the paper.

### Range adaptation is corrected within the decision circuit

As previously shown, *offer value* cells undergo range adaptation (Fig.S3)^23^. Linear tuning implies that any given value interval is allotted the same activity interval in the neuronal representation. Range adaptation ensures that the full activity range is always available to represent the range of values offered in the current context. Thus range adaptation seems to provide an efficient representation for offer values. However, range adaptation also poses a computational puzzle^26^ illustrated in Fig.3ab. In essence, current models assume that binary decisions are made by comparing the firing rates of two neuronal populations encoding the subjective values of the offered goods^27-31^. If so, by varying the ranges of the two offers one could impose any indifference point (an arbitrary choice bias).

**Figure 3.**
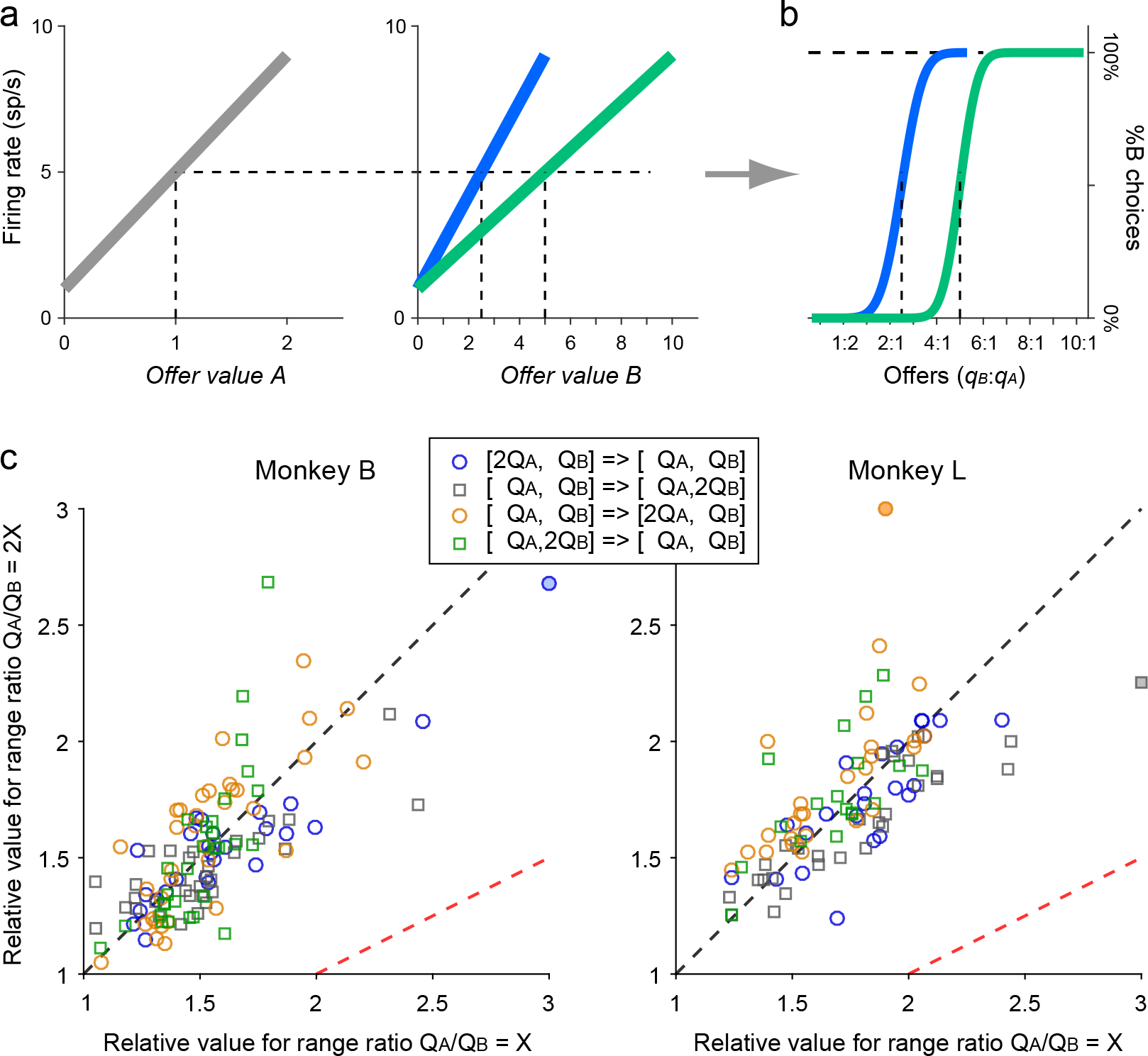
Range adaptation is corrected within the decision circuit. **ab.** Uncorrected range adaptation induces arbitrary choice biases. Panel (a) shows the schematic response functions of two neurons encoding the *offer value* A (left) and the *offer value B* (right). Panel (b) shows the resulting choice patterns under the assumption that decisions are made by comparing the firing rates of these two cells. We consider choices in two conditions, with the range ΔA = [0 2] kept constant. When ΔB = [0 5], the firing rate elicited by offer 1A is between that elicited by offers 2B and 3B (*ρ* = 2.5). When ΔB = [0 10], *offer value B* cells adapt to the new value range. Now offer 1A elicits the same firing rate as offer 5B (*ρ* = 5). Thus if range adaptation is not corrected, changing either value range induces a choice bias. **c.** Relative values measured in Exp.2. The two panels refer to the two animals. In each panel, the axes represent the relative value measured when *Q_A_*/*Q_B_* = X (x-axis) and that measured when *Q_A_*/*Q_B_* = 2X (y-axis). Each data point represents data from one session, and different symbols indicate different protocols (see legend). If decisions were made by comparing uncorrected firing rates, data points would lie along the red dotted line. In contrast, data points lie along the black dotted line (identity line). In other words, the relative values measured in the two trial blocks were generally very similar, indicating that range adaptation was corrected within the decision circuit. Panels ab are reproduced from [^26^].

Exp.2 was conducted to test this prediction in controlled conditions. In each session, monkeys chose between two juices. Trials were divided in two blocks. Across blocks, we either halved or doubled the range of one of the two juices (2x2 design). For each trial block, *Q_A_* and *Q_B_* indicate the maximum quantities of juices A and B offered, respectively. Thus independently of other factors, the ratio *Q_A_*/*Q_B_* changed by a factor of two between blocks (*Q_A_*/*Q_B_* = X or 2X). The experimental design controlled for juice-specific satiety and other possible sources of choice bias (see Methods).

We collected behavioral data in 220 sessions. In each session and each trial block, we measured the relative value of the juices. We then compared the measures obtained in the two trial blocks. According to the argument in Fig.3ab, the relative value measured when *Q_A_*/*Q_B_* = X should be roughly twice that measured when *Q_A_*/*Q_B_* = 2X. Contrary to this prediction, the relative values measured in the two trial blocks were generally similar (Fig.3c). Pooling all sessions, the ratio of relative values measured for the two trial blocks was statistically indistinguishable from 1 (mean ratio = 1.006; p=0.81, Wilcoxon signed rank test) and significantly below 2 (p<10^-37^, Wilcoxon signed rank test). These results held true for each animal.

### Range adaptation maximizes the expected payoff

Exp.2 indicated that range adaptation is corrected within the decision circuit. We previously proposed a possible scheme for this correction. In essence, choice biases are avoided if the synaptic efficacies between *offer value* cells and downstream neuronal populations are rescaled by the value ranges^26,31^. However, if this correction occurs, it is reasonable to question whether range adaptation benefits the decision process at all. The central result of this study is that range adaptation in *offer value* cells maximizes the expected payoff even if adaptation is corrected within the decision circuit. The theoretical argument is summarized here and detailed in the Supplementary Material, where we provide mathematical proofs.

Consider the general problem of choices between two goods, A and B. We indicate the quantities of A and B offered on a particular trial with *q_A_* and *q_B_*. Across trials, *q_A_* varies in the range [0, *Q_A_*], while *q_B_* varies in the range [0, *Q_B_*]. We assume linear indifference curves (Fig.4a) and indicate the relative value with *ρ*. Choices can be described by a sigmoid surface (Fig.4b). For each pair of offers, one of the two options provides a higher payoff, but in some trials the animal fails to choose that option (choice variability). Intuitively, this may happen because the neural decision circuit has a finite number of neurons, limited firing rates, trial-by-trial variability in the activity of each cell, and non-zero noise correlations.

**Figure 4.**
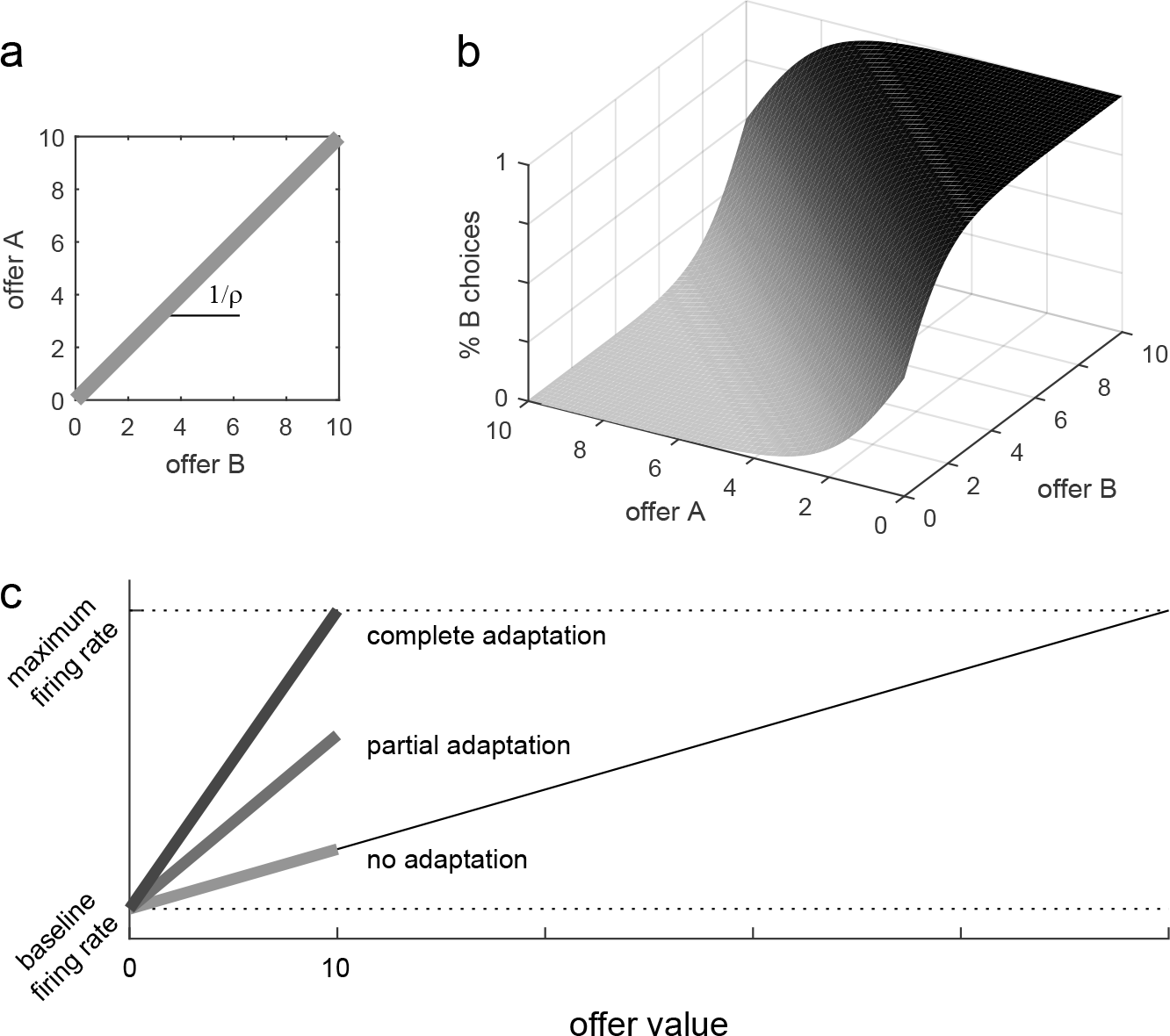
Possible adaptation scenarios. **a.** Indifference line. We indicate with *q_A_* and *q_B_* the quantities of good A and good B, respectively. Across trials, *q_A_* varies in the range [0, *Q_A_*], while *q_B_* varies in the range [0, *Q_B_*]. In the plane defined by *q_A_* and *q_B_*, we define the "indifference curve" as the set of offers for which the animal splits decisions equally between the two goods. We assume that the indifference curve is a straight line on this plane. Thus the relative value between the two goods, referred to as *ρ*, is defined by the slope of the indifference curve (slope = 1/*ρ*). **b.** Choice pattern. Given offers of goods A and B, a choice pattern can be represented as a sigmoid surface, in which the z-axis represents the likelihood of choosing good B. For each pair of offers, one of the two options provides a higher payoff, depending on whether it is above or below the indifference curve. However, unless the sigmoid is a step function, in some trials the animal fails to choose that option (choice variability). **c.** Adaptation scenarios. In this cartoon, offer values in the current context vary in the range [0 10]. The light line represents a hypothetical scenario in which there is no range adaptation (see Results). The darker lines represent the scenarios with partial and complete range adaptation.

Fig.4c illustrates the issue of interest. We assume that neuronal response functions are linear. Actual neurons always have a baseline firing rate (corresponding to a zero offer), but we assume that this activity does not contribute to the decision. Thus we focus on baseline-subtracted response functions. Let us consider a hypothetical scenario in which there is no adaptation. If so, neurons would have fixed tuning, corresponding to a linear response function defined on a very large value range. In contexts where the encoded good varies on a smaller range, neuronal firing rates would span only a subset of their potential activity range. In contrast, if neurons undergo complete range adaptation, firing rates span the full activity range in each behavioral context.

To understand how range adaptation in *offer value* cells affects the expected payoff, it is necessary to consider a specific decision model. That is, the question must be addressed under some hypothesis of how the activity of *offer value* cells is transformed into a decision. We examined the linear decision model^32,33^ formulated as follows:

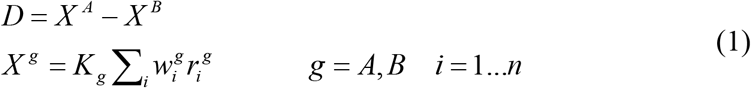

where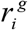 is the firing rate of an *offer value g* cell,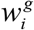 in are decision weights, *n* is the number of cells associated with each juice, and *K_g_* is the synaptic efficacy of *offer value g* cells onto downstream populations. Conditions respectively. *D* > 0 and *D* < 0 correspond to choices of goods A and B, respectively

We model the firing rates of *offer value* cells as Poisson variables and we approximate noise correlations with their mean long-distance component^32^. In accord with experimental measures, we set the noise correlation to *ξ* = 0.01 for pairs of neurons associated with the same good, and to zero for pairs of neurons associated with different goods^32^. Importantly, *ξ* does not depend on firing rates (Fig.S4). We thus compute the probability of choosing juice A given offers *q* = (*q _A_*, *q_B_*), tuning slopes *t* = (*t _A_*, *t _B_*) and synaptic efficacies *K* = (*K _A_*, *K_B_*)

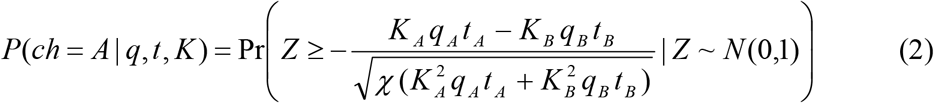

where *N* (0,1) is the standard normal distribution and *χ* = *ξ/* 4

Eq.2, allows to calculate the expected payoff. Indicating with *v* the maximum possible firing rate, we demonstrate that the expected payoff is maximal when *t_g_* = *v/Q_g_*. This condition corresponds to complete range adaptation (Fig.4c). In the symmetric case, defined by *ρ Q_A_* = *Q_B_* (equal value ranges), the expected payoff is maximal when *K_A_ / K_B_* = 1 and there is no choice bias. In the asymmetric case (unequal value ranges), the expected payoff is maximal when *K_A_ / K_B_* ≈ *ρ Q_A_/Q_B_*. In this condition, there is a small choice bias that favors the larger value range and depends on*χ*.

Notably, Eq.2 expresses the sigmoid surface describing choices. By computing the slope of this surface on the indifference line, we show that under optimal coding the steepness of the sigmoid is inversely related to the value ranges (see Supplementary Material, Eq.28).

### Relation between choice variability and value range

The previous section summarizes a theory of optimal coding of offer values for economic decisions. The main prediction for linear response functions is that the slope of the encoding should be inversely proportional to the value range, as indeed observed in the experiments (range adaptation; Fig.S3de). The theory also makes another testable prediction. Consider experiments in which monkeys choose between two juices and value ranges vary from session to session. The sigmoid steepness should decrease as a function of the value ranges. To test this prediction, we examined 164 sessions from Exp.1. For each session, we computed the geometric mean value range Δ ≡ (*ρ Q_A_ Q_B_*)^1/2^, and we obtained a measure for the sigmoid steepness (*η*) from the sigmoid fit. We thus examined the relation between *η* and Δ.

Fig.5ab illustrates the fitted sigmoid obtained for each experimental session in our data set, separately for monkeys V and L. For each animal, sigmoid functions were aligned at the flex and ranked according to Δ. Notably, sigmoid functions with small Δ were generally steeper (large *η*), while sigmoid functions with large Δ were generally shallower (small *η*). In other words, there was a negative correlation between *η* and Δ. This correlation, summarized in a scatter plot (Fig.6), was statistically significant in each animal (monkey V: corr coef = -0.41, p<0.0005; monkey L: corr coef = -0.26, p<0.02). Control analyses confirmed that this result was not due to differences between juice pairings (Fig.S5) or to fluctuations in the relative value (Fig.S6). Similar results were also obtained for data from Exp.2 (Fig.S7).

**Figure 5.**
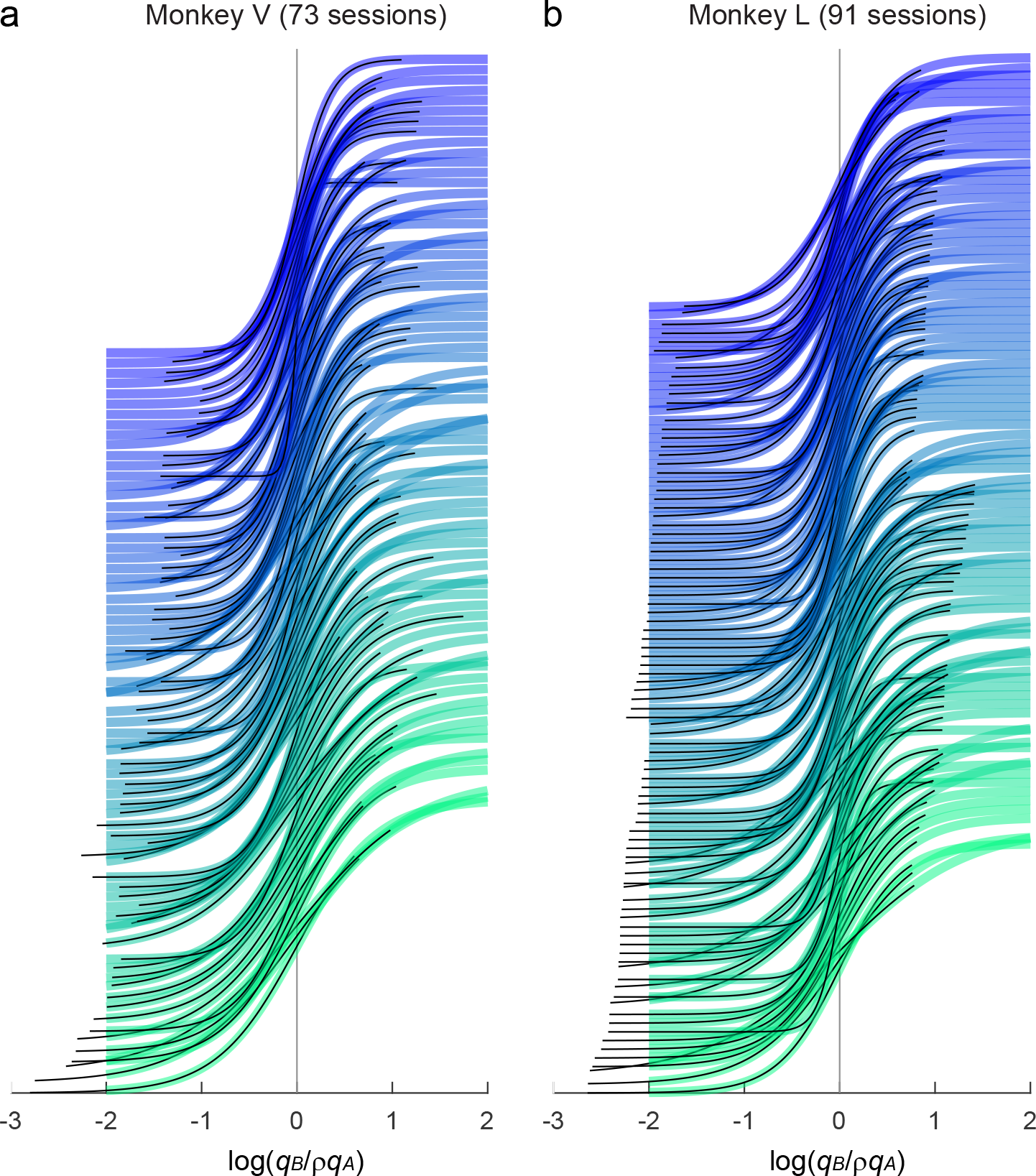
Relation between sigmoid steepness and value range. **a.** Monkey V (73 sessions). For each session, the sigmoid fit provided measures for *ρ* and *η* (Eq.6), and we computed the geometric mean value range Δ. In this plot, different sigmoid functions are aligned at the flex (x-axis) and ranked based on Δ, from bottom (large Δ) to top (small Δ). For each sigmoid, the thick colored line (green-blue) depicts the result of the fit in a standard interval [-2 2]. The thin black line highlights the range of values actually used in the corresponding session. Different shades of color (from blue to green) indicate the ordinal ranking of sessions according to Δ. Notably, sigmoid functions at the bottom of the figure (larger Δ) were shallower (higher choice variability) than sigmoid functions at the top of the figure. **b.** Monkey L (91 sessions). Same format as in (a).

**Figure 6.**
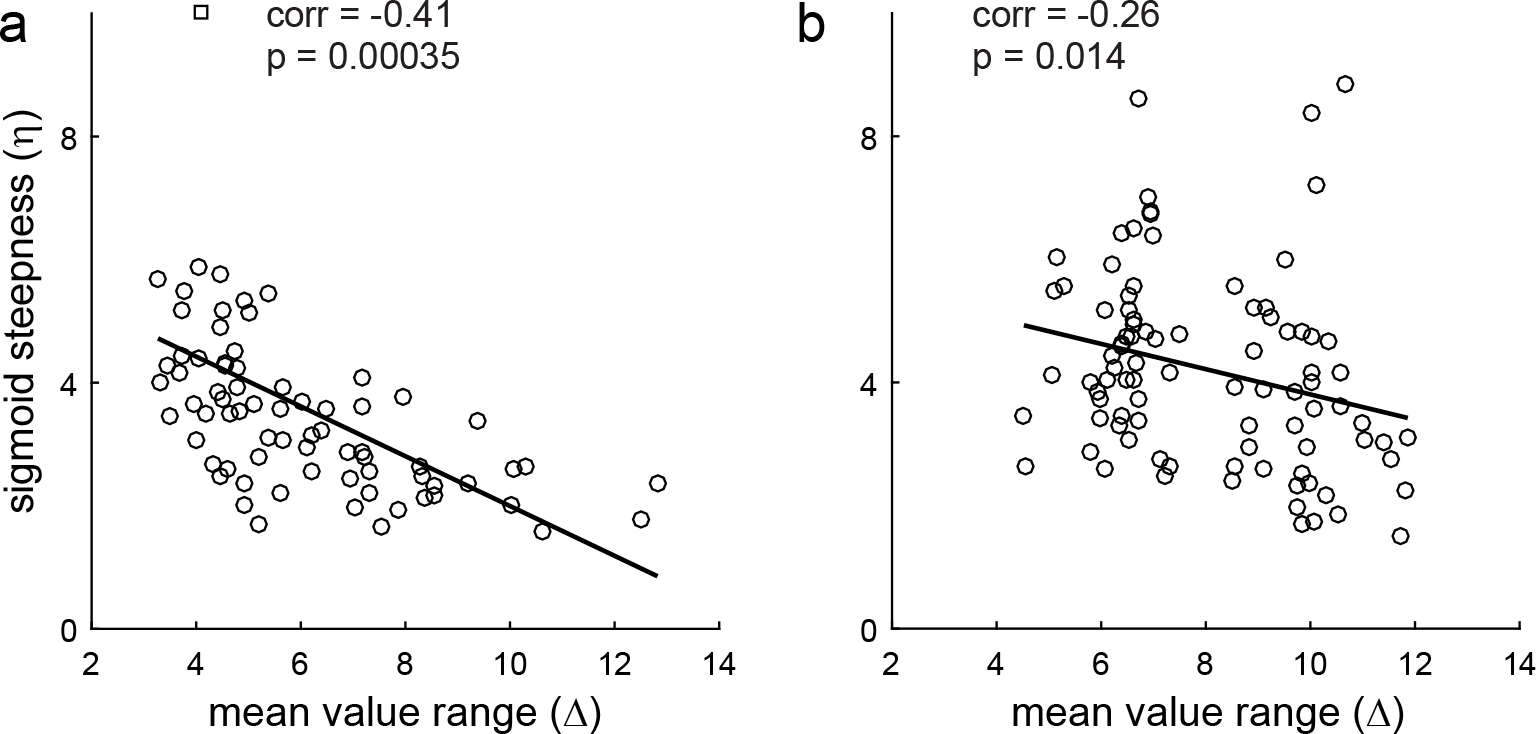
Relation between sigmoid steepness and value range, scatter plots. **ab.** Panels a and b refer to monkey V (73 sessions) and monkey L (91 sessions), respectively. In each panel, the x-axis represents the geometric mean value range Δ ≡ (*ρ Q_A_ Q_B_*)^1/2^, the y-axis represents the steepness of the sigmoid (*η*) and each data point represents one session. In both animals, the two measures were significantly and negatively correlated. In each panel, the black line represents the result of Deming’s regression.

### Neuronal responses are functionally rigid

We have shown that range adaptation maximizes the expected payoff under the assumption of linear response functions. Next we address a closely related question, namely whether (or in what sense) linear response functions are optimal in the first place. In the visual system, optimal coding is achieved if tuning functions match the cumulative distribution of the encoded stimuli^2,7^. In the valuation system, the equivalent condition would occur if *offer value* responses matched the cumulative distribution of offered values. We already showed that this is not the case (Fig.2). In retrospect, this finding is not surprising because a subject performing economic decisions is best served by response functions that maximize the expected payoff, which do not necessarily maximize information transmission. Thus what is the optimal response function for *offer value* cells?

The answer to this question depends on the joint distribution of offers and on the relative value of the two goods. For example, consider the case in which an animal chooses between goods A and B and *ρ* = 2. Good A is always offered in quantity 1, while good B is offered in quantities between 0 and 5 (Fig.7a). We consider *offer value B* cells and we indicate with *r_B_* their firing rate. It is easy to see that the payoff is maximal if *r_B_*(x) = 0 when x<2, *r_B_*(2) = 0.5, and *r_B_*(x) = 1 when x>2, where x are quantities of juice B offered. Hence, the optimal response function is a step function with the step located at x = 2. Next consider the case in which quantities of both goods vary between 0 and 5, at least one of the two goods is always offered in quantity 1, and *ρ* = 2 (Fig.7b). Again, the optimal response function for *offer value B* cells is *r_B_*(x) = 0 when x<2, *r_B_*(2) = 0.5, and *r_B_*(x) = 1 when x>2. For *offer value A* cells, the optimal response function is *r_A_*(0) = 0, *r_A_*(1) = 0.5, and *r_A_*(x) = 1 when x>1. Thus for both goods, the optimal response function is a step function. Analogously, if offer types are the same but *ρ* = 3 (Fig.7c), the optimal response function for *offer value B* cells is a step function with the step located at x = 3.

**Figure 7.**
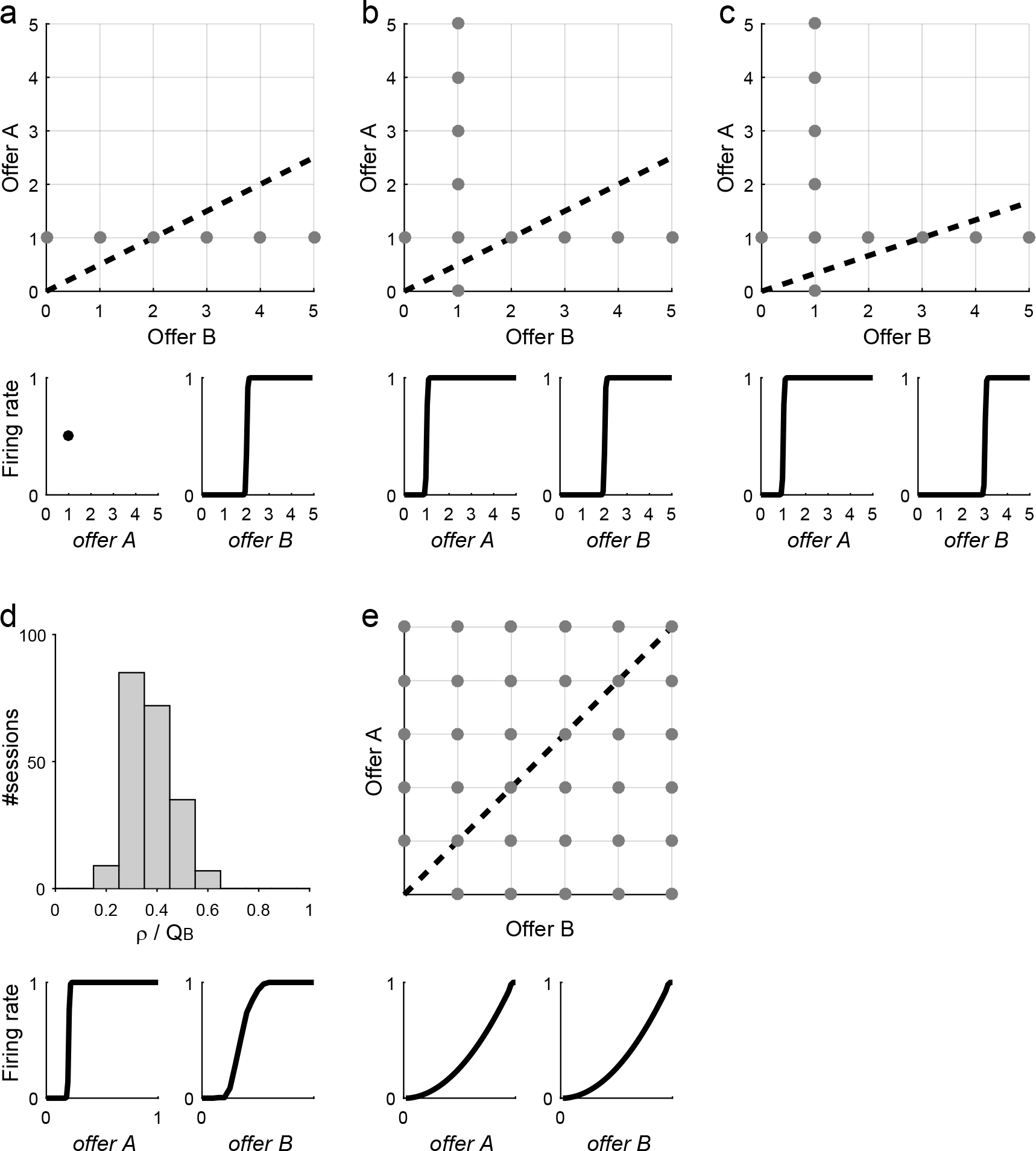
Optimal response functions. **a.** One good offered in fixed quantity (*ρ* = 2). Gray dots represent offer types presented in the session and the dotted line represents the indifference line. Good A is always offered in quantity 1 while good B varies in the range [0 5]. Optimal response functions are shown in the lower panels. **b.** Idealized experimental session (*ρ* = 2). For each good, quantities vary in the range [0 5], but in each offer type at least one good is offered in quantity 1. Lower panels show the optimal response functions (*ORF*, step functions). **c.** Idealized experimental session (*ρ* = 3). **d.** Optimal mean response functions. The histogram represents the distribution of *ρ*/*Q_B_*, where *ρ* is the relative value and *Q_B_* is the maximum quantity of juice B offered. Lower panels show the mean optimal response functions, *mean(ORF)*. For *offer value B*, the response function is computed as the cumulative distribution function for *ρ*/*Q_B_*. **e.** Idealized session with uniform distribution and equal value ranges (a.u.). Lower panels show the corresponding optimal response functions (*ORF_uniform_*). Note that the curvature of *ORF_uniform_* is in the same direction as that observed on average in the neuronal population (Fig.2a, histogram).

The scenarios depicted in Fig.7bc are similar to those occurring in Exp.1. Indeed our sessions always included forced choices for both juices. Furthermore, in 96% (200/208) of our sessions, when both juices were offered, at least one of them was offered in quantity 1 (Fig.S8). Thus in Exp.1, optimal response functions for *offer value* cells would have been step functions, not linear functions. Our neuronal data clearly belied this prediction (Fig.2). In other words, our results indicate that the functional form of *offer value* cells did not adapt to maximize the payoff in each session. To further examine this point, we ran two additional analyses.

First, we entertained the hypothesis that the functional form of *offer value* cells might adapt on a longer time scale, over many sessions. However, we found that the mean optimal response function was a fairly sharp sigmoid (Fig.7d), contrary to our observations (Fig.2). Second and most important, we recognized that neuronal responses examined in Fig.2 were originally identified through a variable selection analysis that only considered linear response functions^21^ (see Methods). This effectively imposed a bias in favor of linearity. To eliminate this bias, we repeated the variable selection procedures including in the analysis all the variables discussed in this study. These included the cumulative distribution function for the number of trials (ntrials_CDF_), the optimal responses in each session (step functions) and the mean optimal response function across sessions (Methods). The results confirmed previous findings: variables *offer value*, *chosen value* and *chosen juice* still provided the highest explanatory power. In particular, the explanatory power of linear offer value variables was significantly higher than that of each of the new variables (Table S1).

In the final analysis of this section, we considered whether the response functions observed experimentally would maximize the expected payoff for other possible joint distributions of offers. One interesting candidate was the symmetric uniform distribution (Fig.7e). We calculated the optimal response functions given this distribution (ORF_uniform_) and we found that they are quasi-linear and slightly convex. Notably, this non-linearity is in the same direction observed in Fig.2a (histogram). We then repeated the variable selection analysis including variables based on ORF_uniform_. Interestingly, neuronal responses best explained by ORF_uniform_ variables were more numerous than those best explained by linear offer value variables (Fig.8). As in previous studies, we used two procedures for variable selection, namely stepwise and best-subset (Methods). Both procedures identified variables *offer A ORF_uniform_*, *offer B ORF_uniform_*, *chosen value* and *chosen juice* as providing the maximum explanatory power (Fig.9, Fig.10). However, a post-hoc analysis indicated that the explanatory power of ORF_uniform_ variables was statistically indistinguishable from that of linear offer value variables (Table S2).

**Figure 8.**
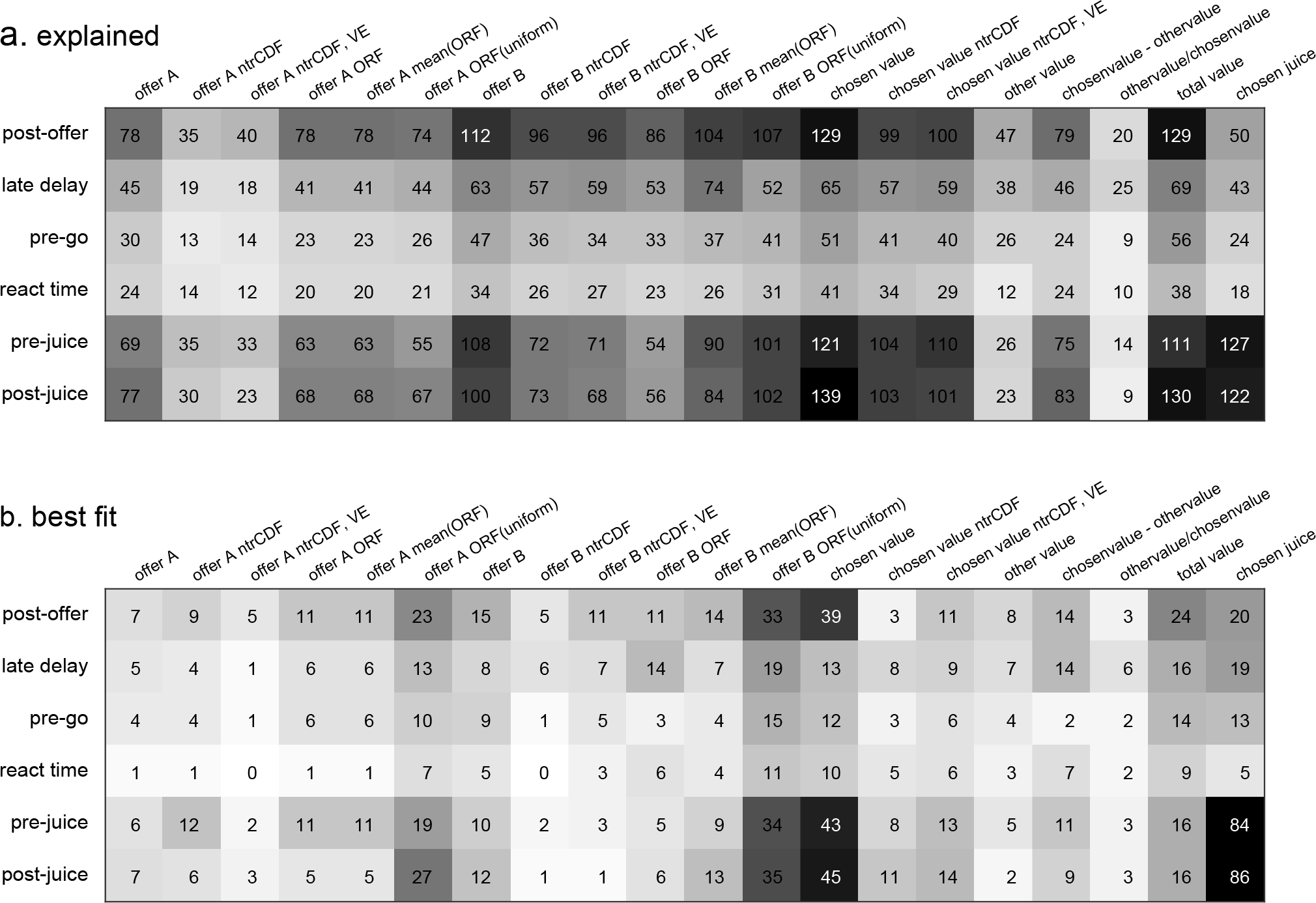
Population analysis of neuronal responses. Each neuronal response that passed an ANOVA criterion was regressed against each variable. If the regression slope differed significantly from zero, the variable was said to explain the response (Methods). **a.** Explained responses. Rows and columns represent, time windows and variables, respectively. In each location, the number indicates the number of responses explained by the corresponding variable. For example, in the post-offer time window, the variable *offer A* (linear response function) explained 78 responses. The same numbers are also represented in grayscale. Each response could be explained by more than one variable. Thus each response might contribute to multiple bins in this panel. **b.** Best fit. In each location, the number indicates the number of responses for which the corresponding variable provided the best fit (highest R^2^). For example, in the post-offer time window, *offer A* (linear response function) provided the best fit for 7 responses. The same numbers are also represented in grayscale. In this panel, each neuronal response contributes at most to one bin. Qualitatively, the dominant variables appear to be *offer A ORF_uniform_*, *offer B ORF_uniform_, chosen value* and *chosen juice*. Indeed the variable selection procedures identified these variables as the ones with the highest explanatory power (Fig.9).

**Figure 9.**
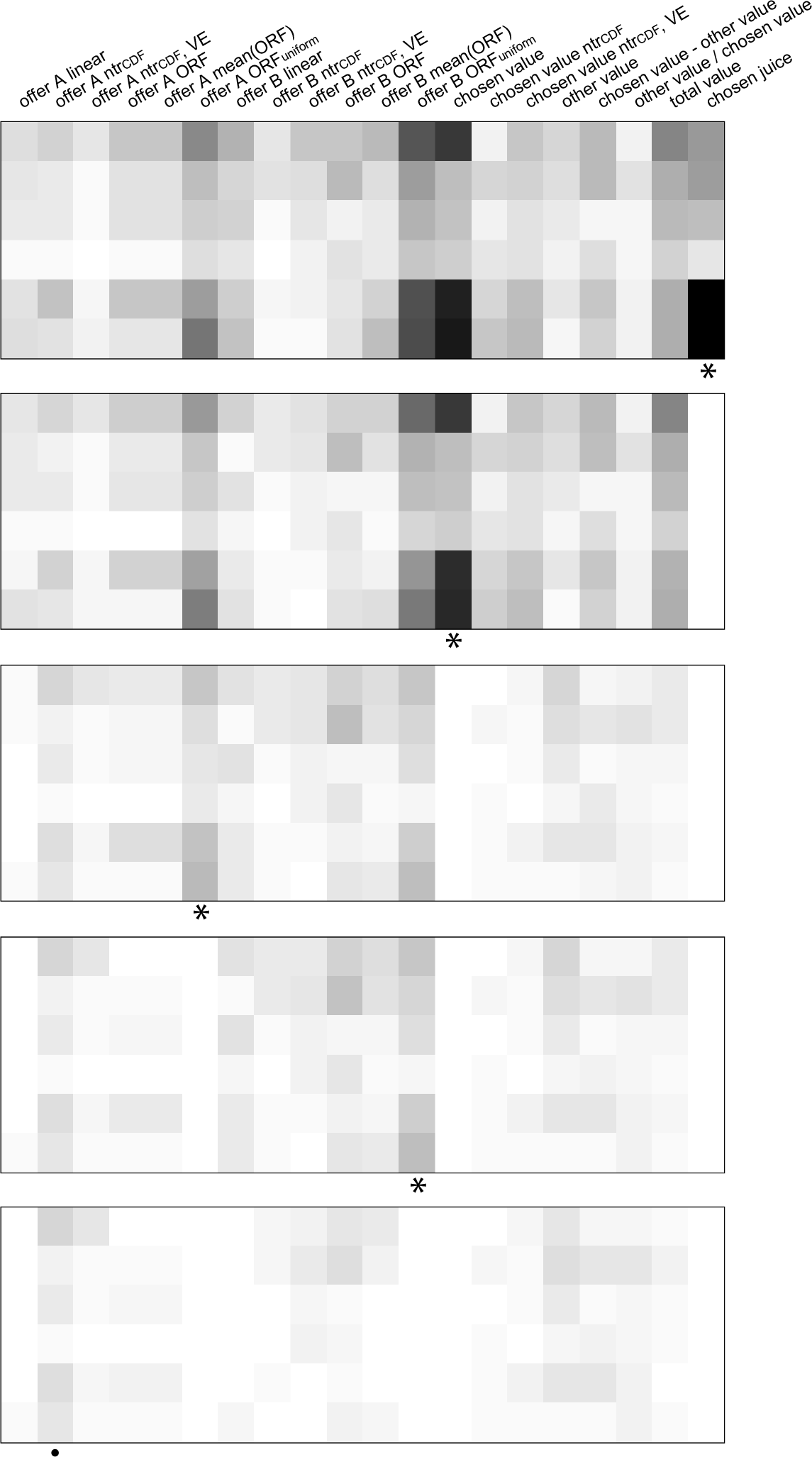
Variable selection analysis, stepwise selection. The top panel is as in Fig.8b. At each iteration, the variable providing the maximum number of best fits in a time window was selected and indicated with a * in the figure. All the responses explained by the selected variable were removed from the pool and the procedure was repeated on the residual data set. Selected variables whose marginal explanatory power was <5% were eliminated (Methods) and indicated with a · in the figure. In the first four iterations, the procedure selected variables *chosen juice*, *chosen value*, *offer A ORF_uniform_* and *offer B ORF_uniform_*, and no other variables were selected in subsequent iterations.

**Figure 10.**
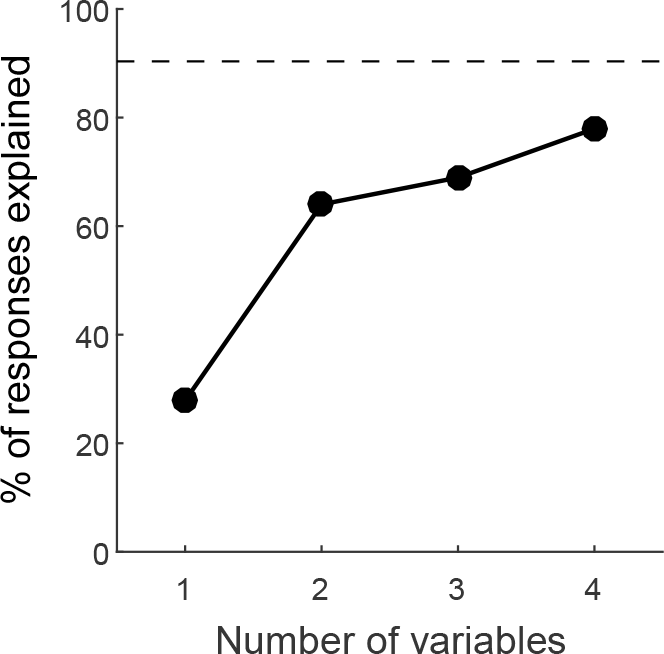
Stepwise selection, percent of explained responses. The y-axis represents the percentage of responses explained at the end of each iteration. The total number of task-related responses (1378) corresponds to 100%. The number of responses explained by at least one of the variables included in the analysis (1245/1378 = 90%) is indicated with a dotted line.

In conclusion, the variable selection analyses confirmed that offer value responses were quasi-linear and thus suboptimal given the joint distributions of offers in our experiments. Furthermore, *offer value* responses were indistinguishable from optimal responses functions calculated assuming a uniform joint distribution of offers. Below we elaborate on the significance of this finding.

### The cost of rigidity and the benefit of adaptation

The tuning of *offer value* cells is functionally rigid (quasi-linear) but parametrically plastic (range adapting with optimal gain). In terms of the expected payoff, functional rigidity ultimately imposes some cost, while range adaptation ultimately yields some benefit. We sought to quantify these two terms in our experiments.

For each session of Exp.1, we focused on strictly binary choices (i.e., we excluded forced choices). Based on the relative value of the juices (*ρ*), we computed for each trial the *chosen value* (i.e., the payoff) and the *max value*, defined as the higher of the two values offered in that trial. We also defined the *chosen value_chance_* as the chosen value expected if the animal chose randomly between the two offers. Hence, *chosen value_chance_* = (*offer value A* + *offer value B*)/2. For each session we defined the *fractional lost value* (FLV) as:

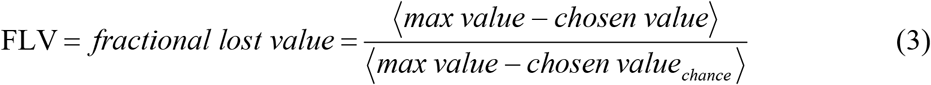

where brackets indicate an average across trials. Under normal circumstances, FLV varies between 0 and 1. Specifically, FLV = 0 if the animal always chooses the higher value (*chosen value* = *max value*) and FLV = 1 if the animal always chooses randomly (*chosen value* = *chosen value_chance_*). Thus FLV quantifies the fraction of value lost to choice variability. For each session, we also computed the *percent error*, defined as the percent of trials in which the animal chose the lower value. We examined these metrics across sessions.

The *percent error* varied substantially from session to session, between 0% and 23% (Fig.S9a). On average across sessions, mean(*percent error*) = 8.7%. The FLV also varied substantially across sessions, between 0 and 0.24 (Fig.S9b). On average across sessions, mean(FLV) = 0.05. Importantly, this estimate provides an upper bound for the value lost by the animal due to suboptimal tuning functions, because other factors might also contribute to choice variability. Hence, the cost of functional rigidity in the coding of offer values may be quantified as ≤ 0.05.

Since we cannot observe decisions in the absence of neuronal adaptation, quantifying the benefits of range adaptation requires a simulation. We proceeded as follows. For each session and for each trial, the sigmoid fit provided the probability that the animal would choose juice B (*P_ch=B_*; see Eq.5) or juice A (*P_ch=A_* = 1 - *P_ch=B_*). Thus in each trial the expected *chosen value* (i.e., the expected payoff) was:

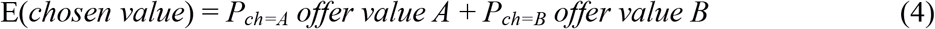

For each session, we computed the *expected fractional lost value* (EFLV) by substituting the E(*chosen value*) for the *chosen value* in Eq.3. Importantly, we verified that EFLV provided a good estimate for the actual FLV (Fig.S9c).

To address the question of interest, we reasoned along the lines of Fig.4c, where the absence of adaptation is approximated with a scenario in which neurons adapt to a very large value range. We already showed that increasing the value range decreases the sigmoid steepness (Fig.6). Thus we examined how reducing the sigmoid steepness would affect the EFLV. We found that the effects were large. For example, when we halved the sigmoid steepness (*η* → *η*/2), we obtained mean(EFLV) = 0.15; when we divided the sigmoid steepness by ten (*η* → *η*/10), we obtained mean(EFLV) = 0.55. Hence, the benefit of range adaptation, while difficult to quantify exactly, is clearly very high.

To summarize, the cost of functional rigidity is rather small compared to the benefit of range adaptation. Our analyses suggest that a quasi-linear but range-adapting coding of offer values is sufficient to ensure close-to-optimal behavior.

## Discussion

Sensory neurons are optimally tuned for perception if they transmit maximal information about the stimuli. In contrast, offer value neurons are optimally tuned for economic decisions if they ensure maximal expected payoff. In this framework, we examined the activity of *offer value* cells in OFC. These neurons are believed to provide the primary input for economic decisions (see below). We showed that their tuning is functionally rigid (linear responses) but parametrically plastic (range adaptation with optimal gain). We also showed that range adaptation is corrected within the decision circuit to avoid arbitrary choice biases. Critically, range adaptation ensures optimal tuning even considering this correction. Confirming theoretical predictions, we showed that choice variability is directly related to the range of values offered in any behavioral context. Finally, we showed that the benefits of range adaptation are large compared to the costs of functional rigidity. Importantly, our theoretical results were derived using a linear decision model (Eq.1)^32,33^. Future work should extend this analysis to other decision models^27,29,31^.

Based on the present results, we cannot confirm whether offer value responses are strictly linear or slightly convex as predicted for optimal response functions under a uniform joint distribution. Thus future work should address this point and consider other joint distributions that might explain neuronal responses in OFC. Nonetheless, the quasi-linear nature of value coding in OFC is noteworthy. We previously showed that the activity of neurons associated to one good does not depend on the identity or value of the other good offered in the same trial^34^. With respect to range adaptation, we also showed that each neuron adapts to its own value range, independently of the range of values offered for the other good^26^. One implication of linear responses (or optimal response functions under a uniform joint distribution) is that the activity of neurons associated with one particular good does not depend on the distribution of values offered for the other good, or on the relative value of the two goods. Thus quasi-linearity can be seen as yet another way in which neurons associated with one particular good are blind to every aspect of the other good. This blindness, termed menu invariance, guarantees preference transitivity^35,36^, which is a fundamental trait of economic behavior. It is tempting to speculate that quasi-linear coding might have been selected in the course of evolution because it facilitates transitive choices.

Adaptive coding has been observed in numerous areas that represent value-related variables including the amygdala^37^, anterior cingulate cortex^38^ and dopamine cells^39-41^. Independently of the specific contribution of each area to behavior, adaptation necessarily poses computational challenges analogous to the coding catastrophe discussed in sensory systems^14,26^. With respect to *offer value* cells in OFC, we previously proposed that choice biases potentially introduced by range adaptation are corrected in the synapses between these neurons and downstream populations^26,31^. The theory of optimal coding developed here makes this same prediction, which should be tested in future experiments. Interestingly, framing^42,43^ and anchoring^44^ effects documented in behavioral economics qualitatively resemble adaptation-driven choice biases, although they are quantitatively more modest. In principle, these effects could be explained if synaptic rescaling trailed neuronal range adaptation. Similar mechanisms have been hypothesized in the visual system to explain illusions and aftereffects^14,17^.

The rationale for this study rests on the assumption that *offer value* cells in OFC provide the primary input for the neural circuit that generates economic decisions. Support for this assumption comes from lesion studies^45-47^, from the joint analysis of choice probability and noise correlation^32^ and from the relation between choice variability and value range shown here. Indeed, current neuro-computational models of economic decisions embrace this view^28,31,48-51^. However, causal links between the activity of *offer value* cells and the decision have not yet been demonstrated with the gold-standard approach of biasing choices using electrical or optical stimulation. Future work should fill this important gap.

## Code availability

The code used for data analysis and simulations is available from the corresponding author upon reasonable request.

## Data availability

The data that support the findings of this study are available from the corresponding author upon reasonable request.

## Acknowledgements

This work was partly conducted while C.P.-S. was a visiting fellow at the Italian Institute of Technology. We thank Heide Schoknecht for help with animal training, Nicolas Brunel and Daniel Chicharro for helpful conversations, and Harold Burton, Arno Onken and Stefano Panzeri for comments on the manuscript. This work was supported by the National Institutes of Health (grant numbers R01-DA032758 and R01-MH104494 to C.P.-S. and grant numbers T32-GM008151 and F31-MH107111 to K.E.C.) and by the National Science Foundation (grant SES-1357877 to A.R.).

## Author Contributions

A.R. and C.P.-S. designed the study; K.E.C, X.C. and C.P.-S. collected the experimental data; K.E.C. and C.P.-S. analyzed the experimental data; A.R. and C.P.-S. developed the mathematical formalism; A.R. and C. P.-S. wrote the manuscript.

---

**Figure S1.**
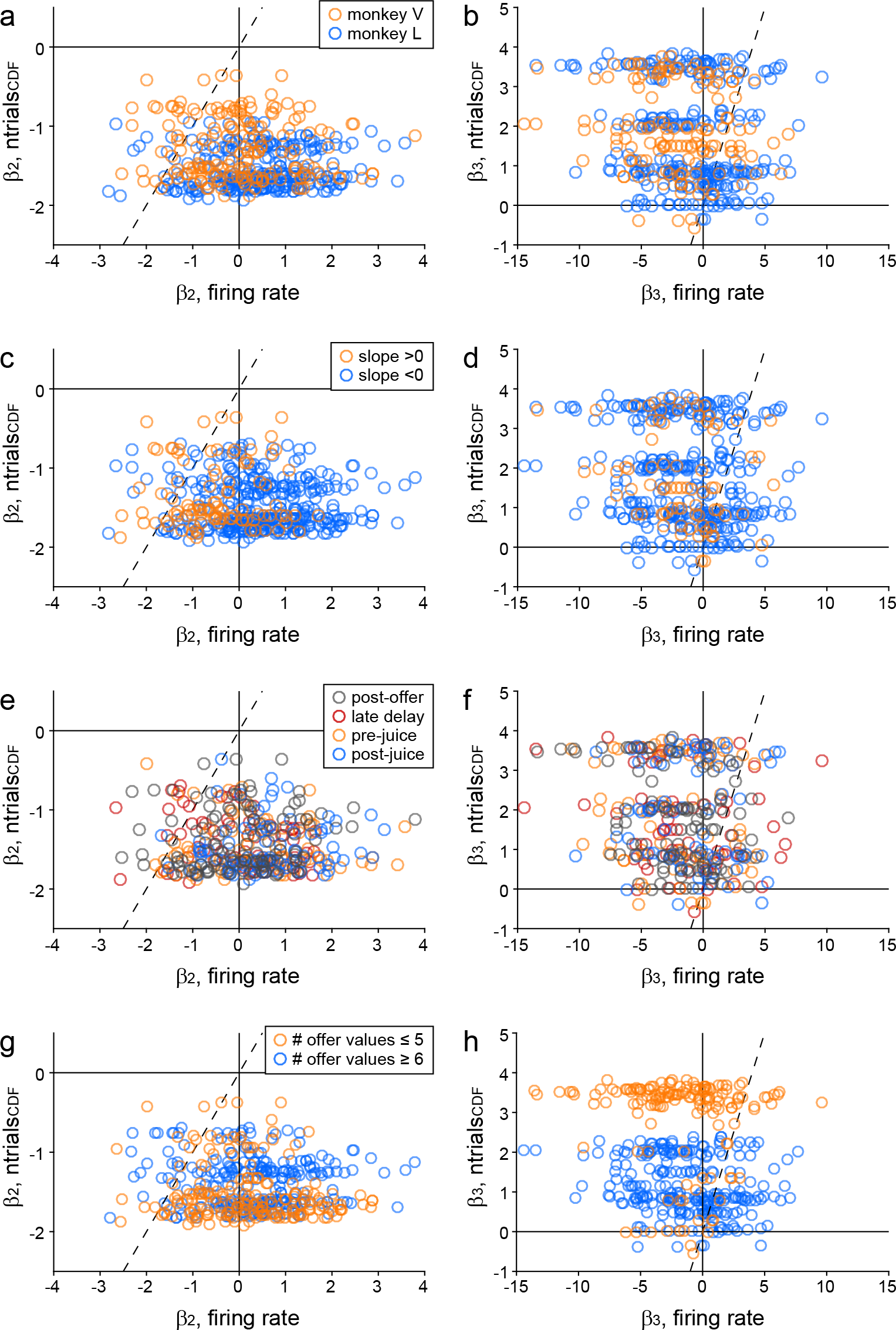
Linearity of neuronal responses, control analyses. **ab.** Individual monkeys. Both panels illustrate the same data shown in Fig.2ab, and different colors indicate neuronal responses from the two monkeys. **cd.** Sign of the encoding. The neuronal encoding of value could be positive (higher firing rates for higher values, or slope>0) or negative (higher firing rates for lower values, or slope<0). In this case, different colors indicate neuronal responses with the two slope signs. **ef.** Individual time windows. In Fig.2ab, we pooled data from different time windows. Here different colors indicate responses from individual time windows. For clarity, we included in panels (ef) only responses from the four primary time windows. Thus some data points present in the other panels are missing here. **gh.** Number of offer values. The number of offer values varied for different responses, and was typically lower for *offer value A* than for *offer value B* responses (see Fig.1). Here different colors indicate that the number of value offered was low (≤5) or high (≥6).

**Figure S2.**
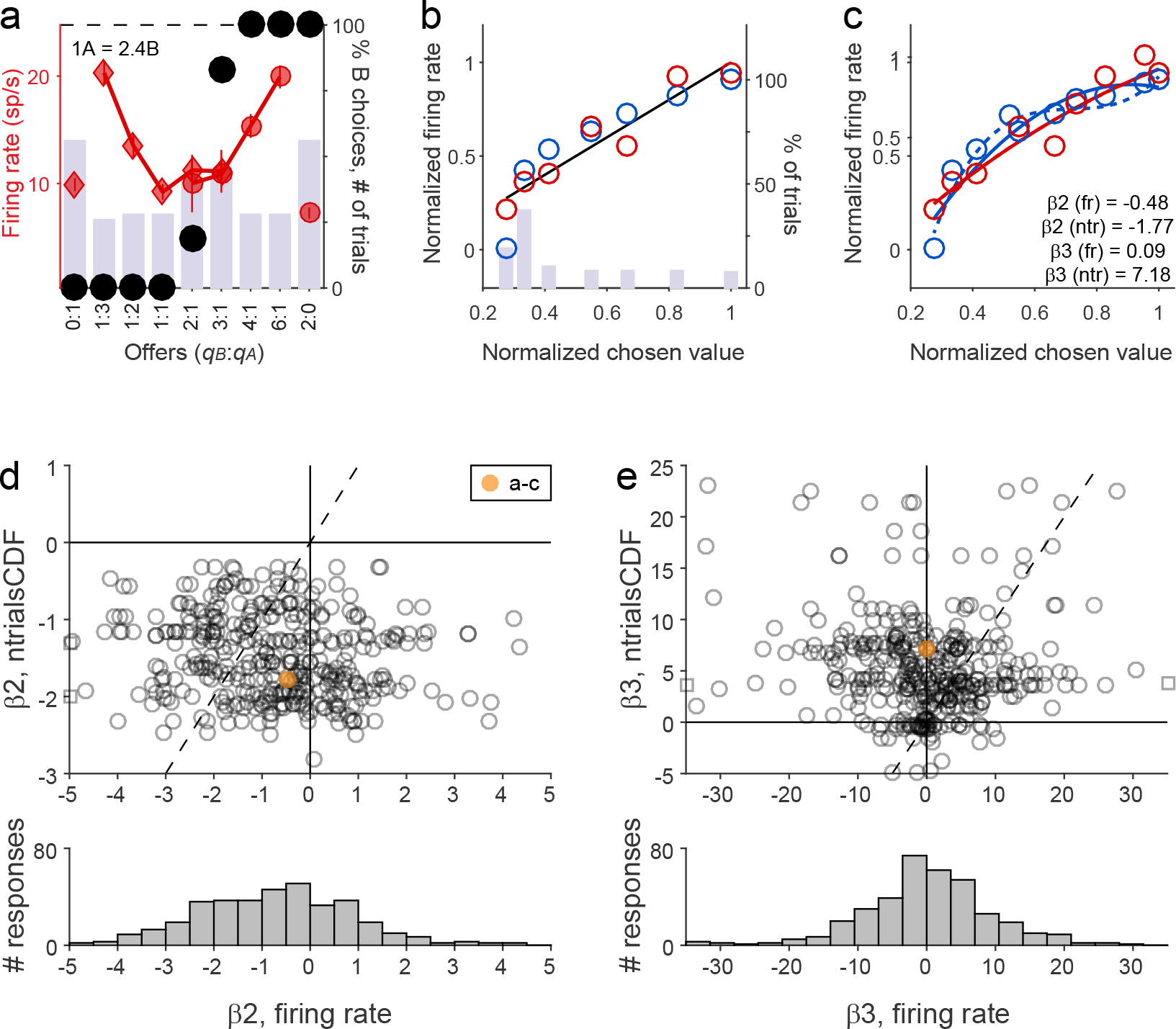
Linearity of *chosen value* responses. **a-d.** Example. Same format as in Fig.2. **ef.** Population analysis (N = 370). Same format as in Fig.2.

**Figure S3.**
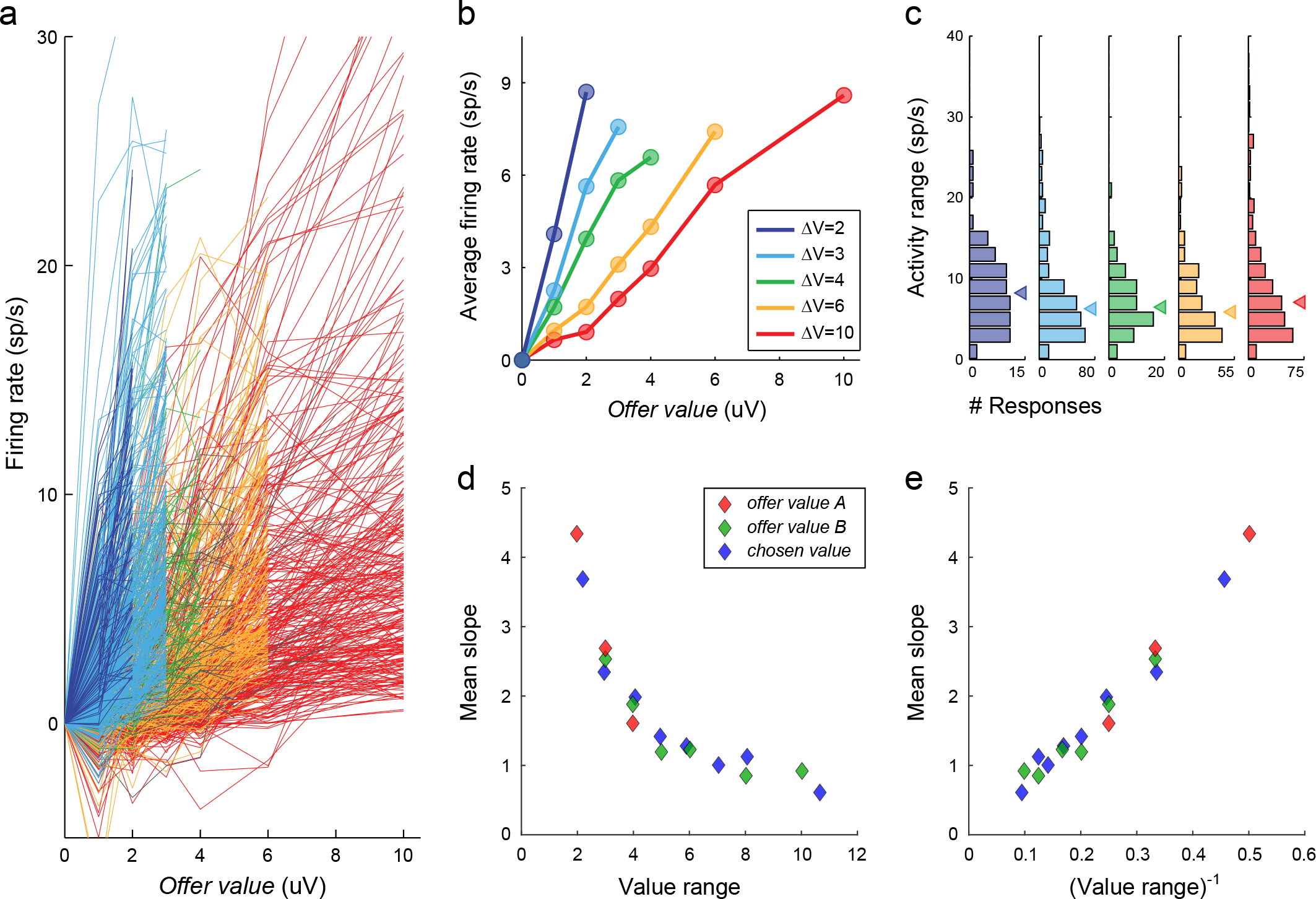
Range adaptation in OFC. **ab.** Individual responses and population averages. Each line in (a) represents one neuronal response. Responses were baseline-subtracted and color-coded according to the range of values offered for the encoded juice. Responses with positive encoding (increasing firing rates for increasing values) and negative encoding (decreasing firing rate for increasing values) were pooled. Each color group presented a wide distribution of firing rates. However, range adaptation became clear once responses were averaged different within each group (panel b). **c.** Distribution of activity ranges. Each histogram illustrates the distribution of activity ranges recorded with the different value ranges. **de.** Mean tuning slope. In (d), each symbol represents the tuning slope (y-axis) averaged across all the neuronal responses recorded with a given value range (x-axis). The three colors indicate different groups of cells (see legend). In (e), the same data points are plotted against the inverse value range (x-axis). Reproduced from [^23^].

**Figure S4.**
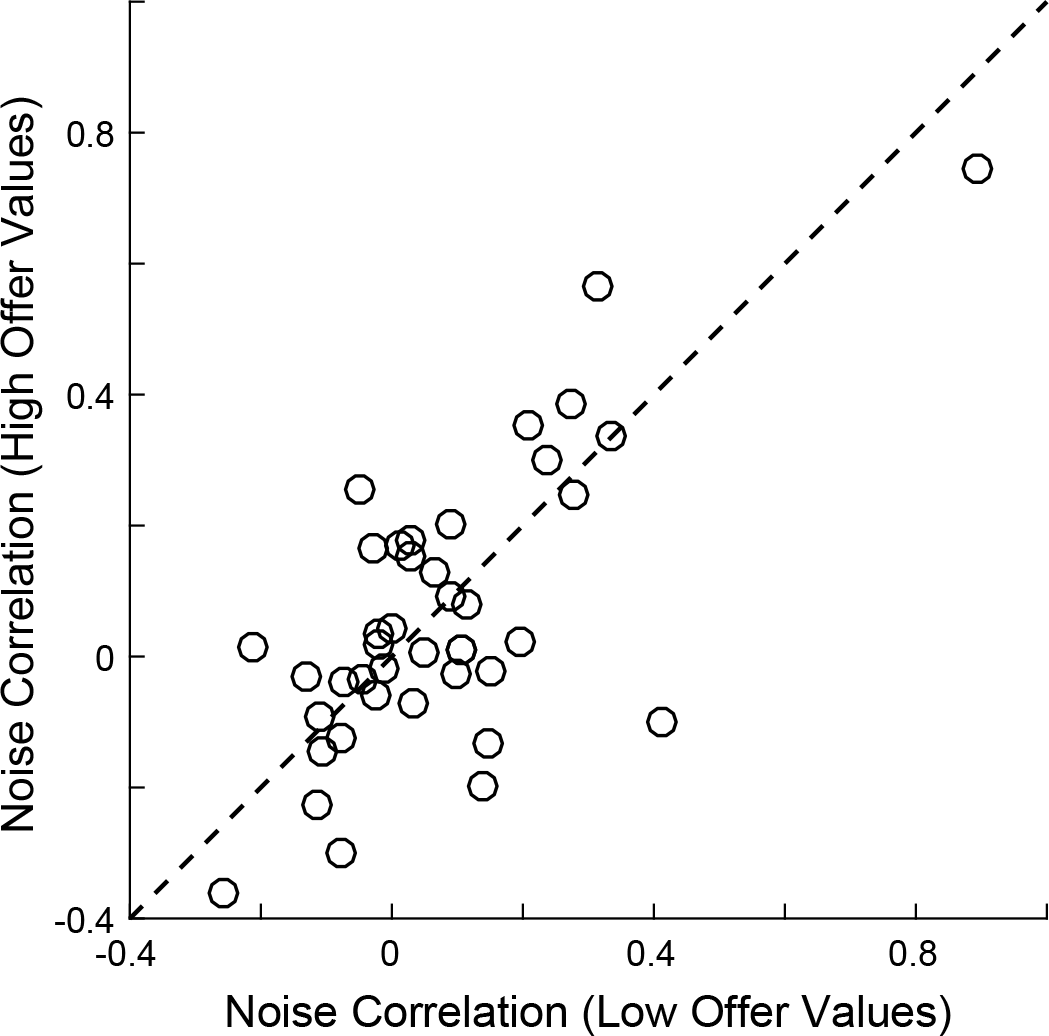
The theoretical argument showing that range adaptation ensures maximum payoff assumes that noise correlation (*ξ* in Eq.2) does not depend on the slope of the encoding. In other words, we assumed that range adaptation would not affect noise correlation. Noise correlation in Exp.1 was analyzed in a previous study, where we found a weak but significant relation between the baseline firing rates and noise correlation (Fig.4c in [^32^]). In other words, pairs of neurons with higher firing rates were slightly more correlated. However, this finding is not directly relevant to the issue of interest here, namely whether noise correlation depends on the firing rates given a pair of cells. To address this issue, we re-examined the same data set focusing on pairs of *offer value* cells recorded simultaneously, associated with the same juice and with the same coding sign (N = 41 pairs; see [^32^] for detail). For each cell pair, we divided trials based on whether the offer value was above or below the median offer value in that session. (Trials in which the value was exactly equal to the median were excluded.) We then computed the noise correlation separately for the two groups of trials. The scatter plot illustrates the results obtained for this population. Each data point represents one cell pair, and the x- and y-axes represent the noise correlation measured in low-value trials (*ξ_low_*) and in high-value trials (*ξ_high_*), respectively. Notably, noise correlation did not differ systematically between the two groups of trials (median(*ξ_high_* -*ξ_low_*) = 0.001; p = 0.62, paired t-test). Similar results were obtained by inverting the axes for pairs of cells with negative encoding (median(*ξ_high_* -*ξ_low_*) = 0.006; p = 0.47, paired t-test) any by excluding pairs of cells from the same electrode (median(*ξ_high_* -*ξ_low_*) = 0.003; p = 0.33, paired t-test).

**Figure S5.**
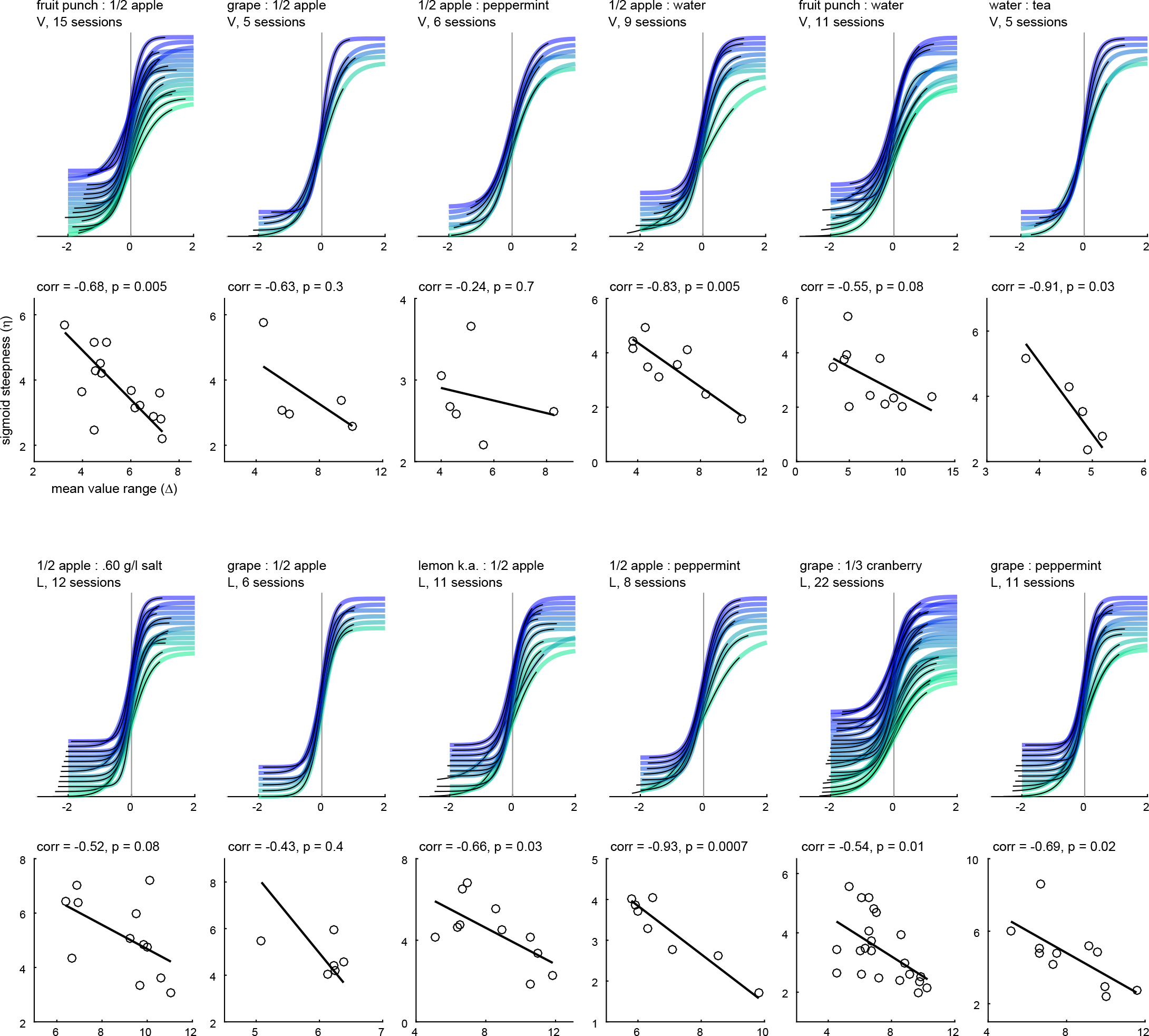
Relation between sigmoid steepness (*η*) and value range (Δ), by juice pair. Fig.6 includes sessions with different juice pairs, with different typical values for *ρ*. In principle, choice variability could vary from juice pair to juice pair in a way that induces the relation between *η* and Δ. To address this issue, we divided sessions in different sets based on the juice pair. Considering only sets with ≥5 sessions, our data included 12 viable sets (6 from each monkey). Here we illustrate the analysis restricted to individual data sets. For each set, the top panel illustrates the fitted sigmoids (equivalent to Fig.5) and the bottom panel illustrates the relation between *η* and Δ (equivalent to Fig.6). For each set we indicate explicitly the juice pair, the animal (V or L) and the number of sessions available. In the bottom panel, we also indicate the correlation coefficient with its p value (as in Fig.6). Black lines illustrate the result of Deming’s regressions. Notably, the negative correlation between *η* and Δ can be observed for each set.

**Figure S6.**
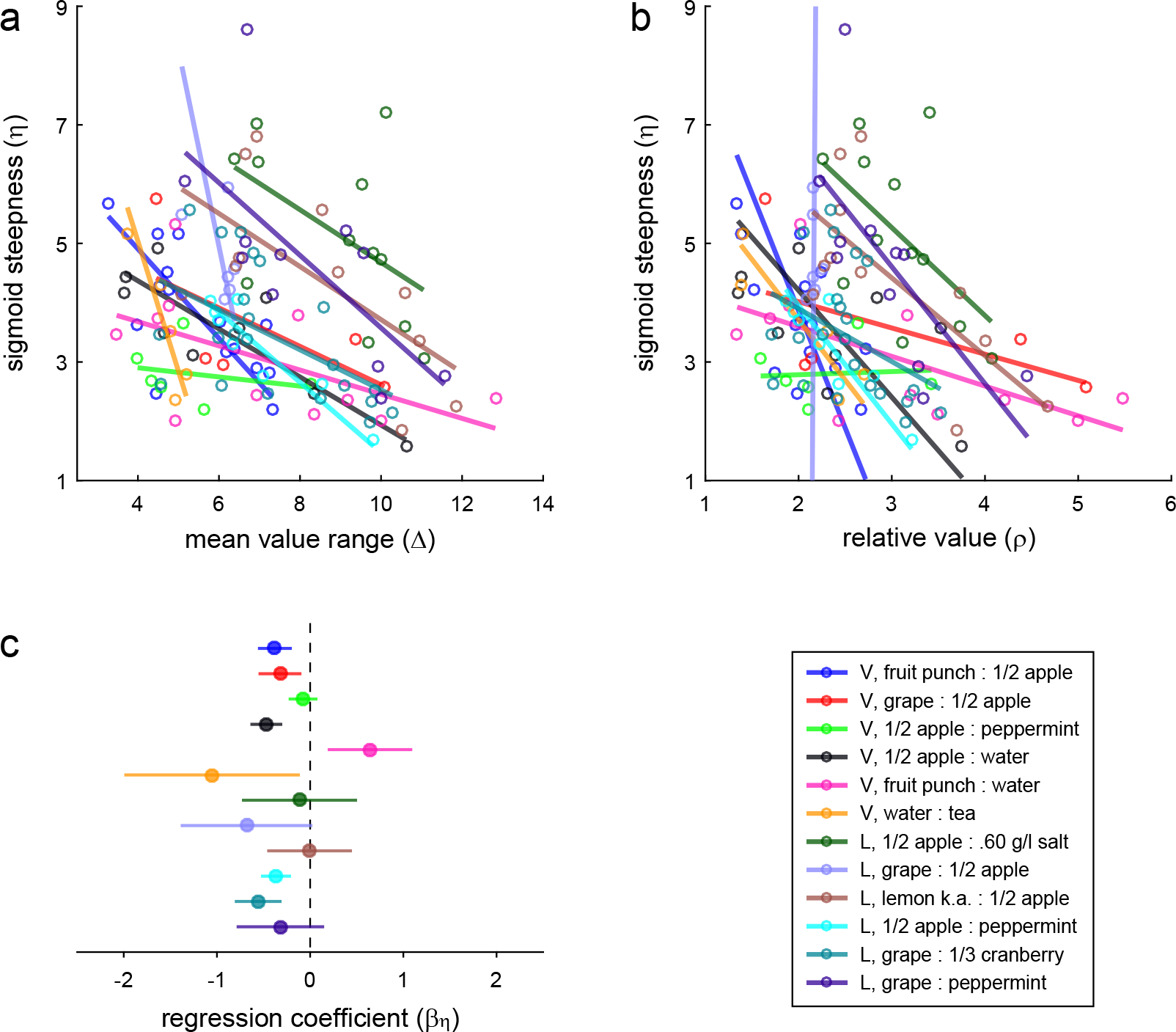
Relation between sigmoid steepness (*η*) and value range (Δ), controlling for fluctuations in relative value (*ρ*). Because *η*, *ρ* and Δ are inter-related (see Methods), the relation between *η* and Δ (Fig.6) might simply reflect fluctuations in *ρ*. To address this issue, we regressed *η* on *ρ* and then on Δ in a stepwise way. The coefficient obtained from the second regression (*β_η_*) quantified the correlation between *η* and Δ not explained by fluctuations of *ρ*. **a.** Relation between sigmoid steepness (*η*) and value range (Δ). This panel recapitulates Fig.S5. Different colors indicate different juice pairs (see legend). **b.** Relation between sigmoid steepness (*η*) and relative value (*ρ*). **c.** Results of stepwise regression. For most sets, we found *β_η_* < 0, indicating that the residual *η* after regressing on *ρ* was still negatively correlated with Δ. Error bars represent S.E.M. Considering the 12 sets, the distribution of *β_η_* was significantly displaced from zero (mean(*β_η_*) = -0.31, p = 0.01, one-tailed t test). In other words, the negative correlation between *η* and Δ was above and beyond the correlation explained by fluctuations of *ρ*. Color conventions in (b) and (c) are as in (a).

**Figure S7.**
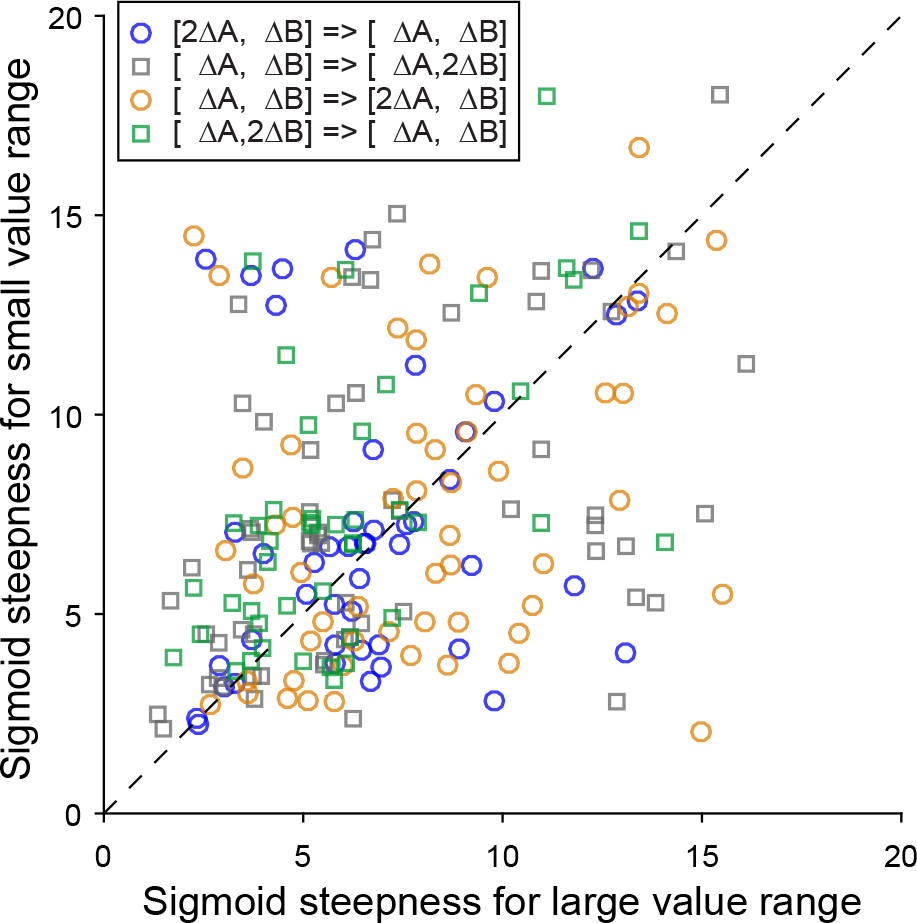
Steepness of the sigmoids measured in Exp.2. For each session, the two axes represent the steepness of the sigmoid measured with a large value range (x-axis) and that measured with a small value range (y-axis). Each data point represents one session. Different symbols represents the block order (see legend) and data from the two monkeys are pooled. Data points are broadly scattered, but overall they tend to lie above the identity line (p = 0.04, sign test). In other words, the sigmoid was steeper when value ranges were smaller. This effect was statistically significant when the analysis was restricted to sessions in which we varied the range of juice B (p = 1.5 10^-4^, sign test), but not when the analysis was restricted to sessions in which we varies the range of juice A (p = 0.37, sign test). Sessions that presented perfect separation (saturated choice patterns) were excluded from this analysis.

**Figure S8.**
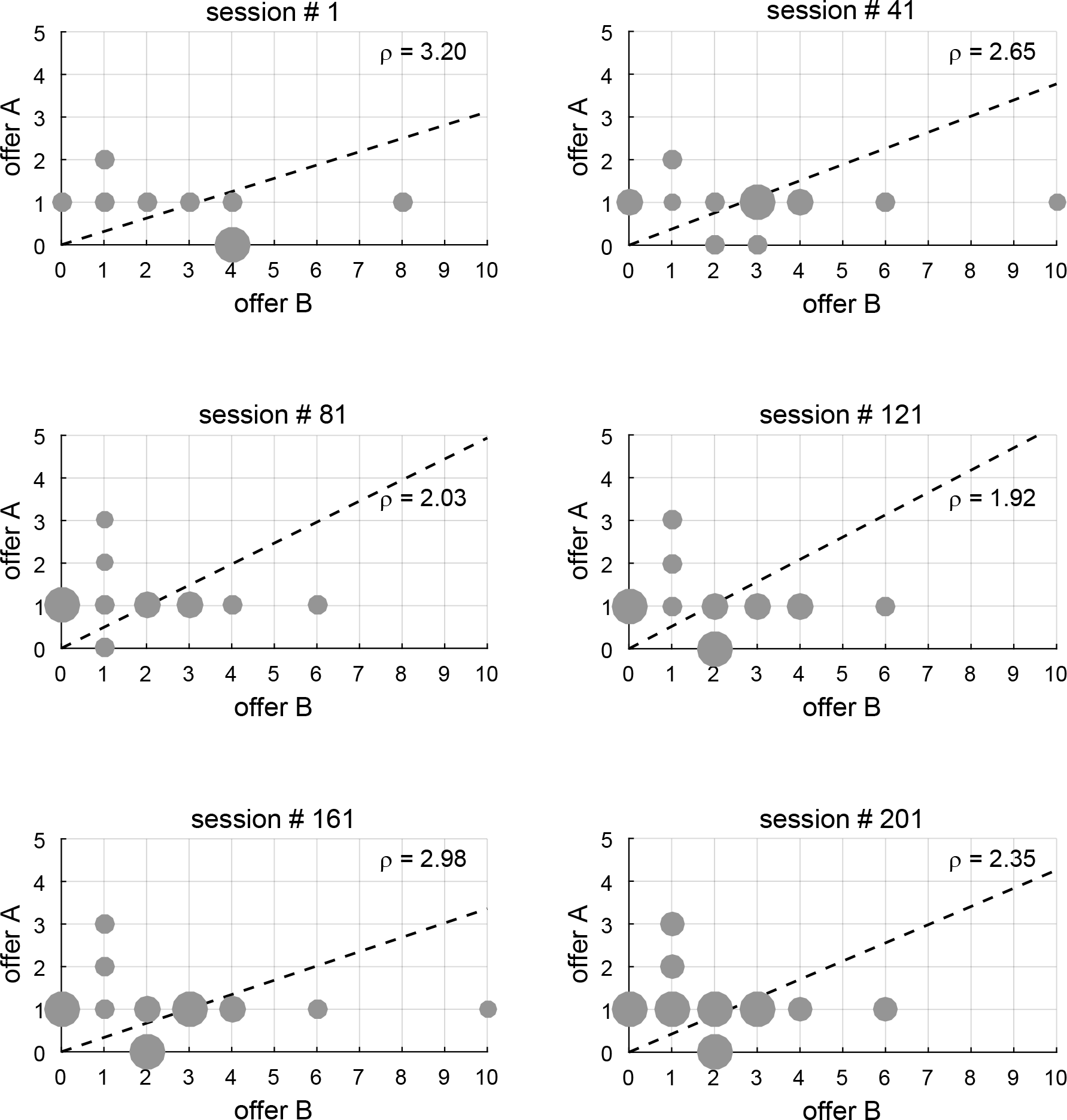
Joint distributions of offers in Exp.1. The six panels illustrate the joint distribution of offers for six representative sessions (out of 208). For each session, gray dots represent offer types presented in the session and the dotted line represents the indifference line (*ρ* calculated with the sigmoid fit). The radius of each dot is proportional to *#trials*/*max(#trials)*, where *#trials* is the number of trials for the corresponding offer type and *max(#trials)* is the maximum of *#trials* across offer types. For the six sessions, *max(#trials)* was equal to 1,429 (session 1), 441 (session 41), 347 (session 81), 400 (session 121), 362 (session 161), and 228 (session 201). Note that the range of offer values and the relative multiplicity of trials for different offer types varied from session to session. Each session included forced choices for either juice. In the other trials, one of the two juices was always offered in quantity 1.

**Figure S9.**
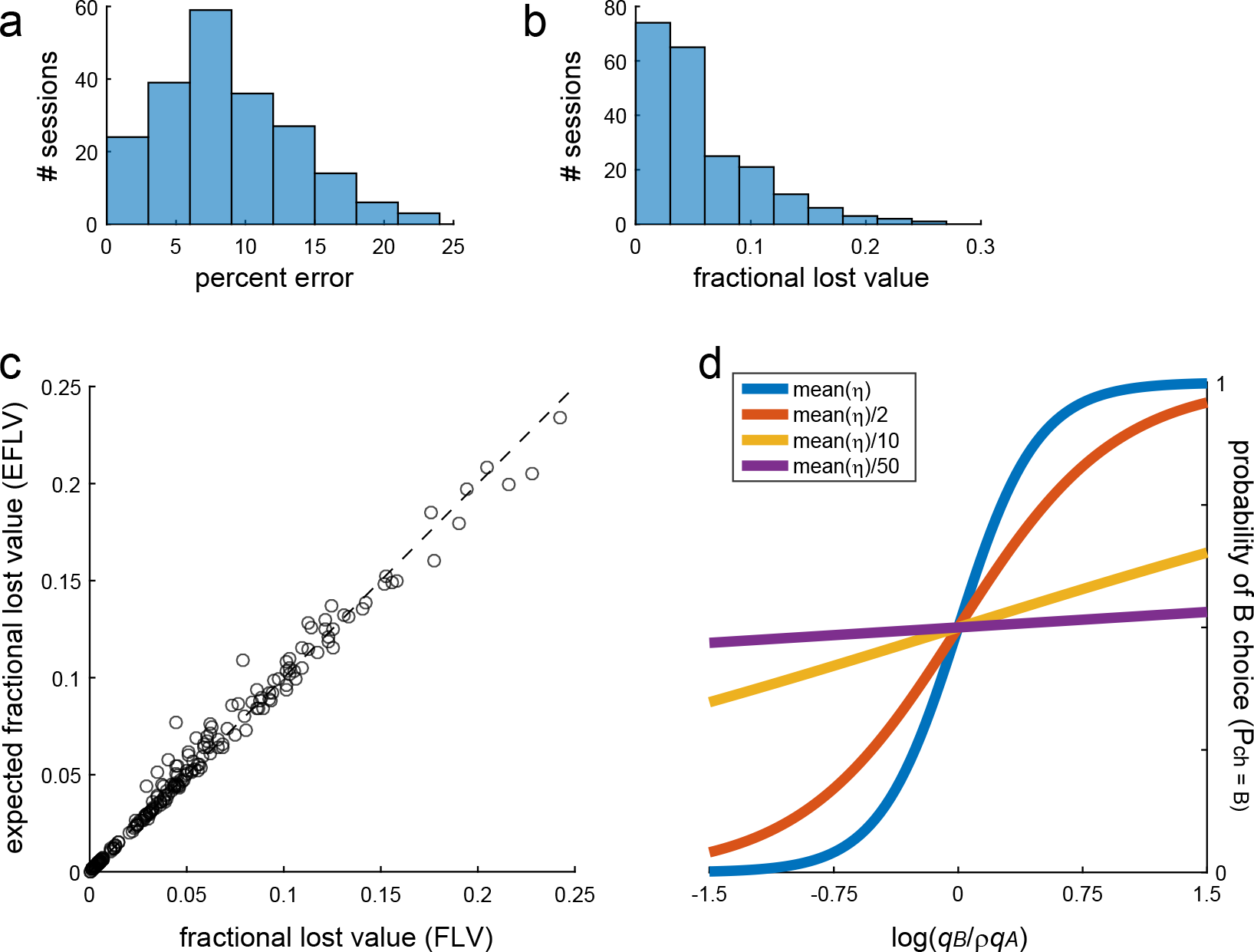
Analysis of lost value. **a.** Distribution of *percent errors* across sessions. On average across sessions, mean(*percent errors*) = 8.7%. **b.** Distribution of *fractional lost value* across sessions. On average across sessions, mean(FLV) = 0.054. **c.** Expected versus actual *fractional lost value*. Each data point represents one session. The *expected fractional lost value* (EFLV; y-axis) was almost identical to the *fractional lost value* (FLV; x-axis). On average across sessions, mean(EFLV) = 0.055. **d.** Effects of decreasing the steepness of the sigmoid. The range on the x-axis is realistic for our experiments. The blue line is obtained with the mean sigmoid steepness measured in Exp.1 (mean(*η*)). The red, yellow and purple lines were obtained by dividing mean(*η*) by 2, 10 and 50, respectively.

**Table S1.** Variable selection, post-hoc analysis. The variable selection analyses were conducted twice. This table refers to the analysis that did not include variables based on *ORF_uniform_* (Methods). Both stepwise and best-subset methods selected variables *offer A linear*, *offer B linear*, *chosen value* and *chosen juice*. The table indicates the results of the post-hoc analysis based on collapsed variables (Methods). To test whether the explanatory power of *offer A|B linear* (column X) was statistically higher than that of other variables examined in the analysis (column Y), we calculated the marginal explanatory power of each variable. For example, considering variables X and Y, the marginal explanatory power of X (column R^2^X) was defined as the total R^2^ explained by X but not by Y. The two variables were then compared with a binomial test. The results (p value in the right column) indicate that the explanatory power of linear offer value variables was significantly higher than that of any competing variable.

| Variable X | Variable Y | R<sup>2</sup>X | R<sup>2</sup>Y | pval |
| --- | --- | --- | --- | --- |
| *offer A\|B linear* | *offer A\|B ntrial<sub>CDF</sub>* | 88.8 | 27.4 | <10<sup>-8</sup> |
| *offer A\|B linear* | *offer A\|B ntrial<sub>CDF</sub> VE* | 86.1 | 27.8 | <10<sup>-8</sup> |
| *offer A\|B linear* | *offer A\|B ORF* | 69.6 | 29.2 | <10<sup>-5</sup> |
| *offer A\|B linear* | *offer A\|B mean(ORF)* | 48.0 | 27.5 | 0.009 |

**Table S2.** Variable selection, post-hoc analysis. This table refers to the analysis that included all the variables, including those based on *ORF_uniform_*. Both stepwise and best-subset methods selected variables *offer A ORF_uniform_*, *offer B ORF_uniform_*, *chosen value* and *chosen juice* (see Fig.9). Same format as Table S1. Note that all comparisons are statistically significant except that between *offer A|B ORF_uniform_* and *offer A|B linear*.

| Variable X | Variable Y | R<sup>2</sup>X | R<sup>2</sup>Y | pval |
| --- | --- | --- | --- | --- |
| *offer A\|B ORF<sub>uniform</sub>* | *offer A\|B linear* | 37.7 | 32.5 | 0.270 |
| *offer A\|B ORF<sub>uniform</sub>* | *offer A\|B ntrial<sub>CDF</sub>* | 114.3 | 47.8 | <10<sup>-7</sup> |
| *offer A\|B ORF<sub>uniform</sub>* | *offer A\|B ntrial<sub>CDF</sub> VE* | 111.4 | 48.0 | <10<sup>-7</sup> |
| *offer A\|B ORF<sub>uniform</sub>* | *offer A\|B ORF* | 93.2 | 47.7 | <10<sup>-4</sup> |
| *offer A\|B ORF<sub>uniform</sub>* | *offer A\|B mean(ORF)* | 78.0 | 52.5 | 0.013 |

## Methods

### Experimental procedures

All experimental procedures conformed to the NIH *Guide for the Care and Use of Laboratory Animals* and were approved by the Institutional Animal Care and Use Committees at Harvard University (Exp.1) and Washington University in St Louis (Exp.2). No subject randomization or blinding during data analysis was used.

The procedures for Exp.1 have been previously described^21^. Briefly, one male (V, 9.5 kg) and one female (L, 6.5 kg) rhesus monkey participated in the experiment. Animals sat in an electrically insulated enclosure with the head restrained, and a computer monitor was placed in front of them at 57 cm distance. In each session, the monkey chose between two juices, labeled A and B, with A preferred. The range of quantities offered for each juice remained fixed within a session, while the quantity offered on any given trial varied pseudo-randomly. Across sessions, we used various quantity ranges for the two juices. The minimum quantity was always zero drops (forced choice for the other juice), while the maximum quantity varied from session to session between 2 and 10 drops. At the beginning of each trial, the animal fixated a center position on the monitor (Fig.1a). After 0.5 s, two sets of colored squares appeared on the two sides of the center fixation. The two sets of squares represented the two offers, with the color associated with a particular juice type and the number of squares indicating the juice quantity. The animal maintained center fixation for a randomly variable delay (1-2 s), at the end of which the center fixation point was extinguished (go signal). The animal revealed its choice by making a saccade towards one of two targets located by the offers, and maintained peripheral fixation for an additional 0.75 s before the chosen juice was delivered. While animals performed in the task, we recorded the activity of individual neurons from the central OFC (see below).

In Exp.2, animals performed essentially the same task, except that sessions were divided into two blocks of trials. One male (B, 9.0 kg) and one female (L, 6.5 kg) rhesus monkey participated in the experiment. The task was controlled through custom-written software based on Matlab (MathWorks)^52,53^ and gaze direction was monitored with an infrared video camera (Eyelink, SR research). The trial structure was the same as Exp.1, except that the initial fixation lasted 1.5 s. Each session included two trial blocks. The minimum offered quantity for each juice was always set to zero (forced choice for the other juice). The maximum quantity (and thus the range) varied from session to session and from block to block. In the second block, we either halved or doubled the range for one juice (A or B) while keeping the other range unchanged. This procedure resulted in a 2x2 design. Each block included 110-260 trials. In each block, an "offer type" was defined by a pair of offers (e.g., [1A:3B]); a "trial type" was defined by an offer type and a choice (e.g., [1A:3B, B]). The relative value of the two juices was computed from the indifference point (see below).

In principle, changes in relative value could arise from factors other than the value range. Exp.2 was designed to minimize three potential sources of choice bias. First, in previous work, we often noted that the relative value of any two juices tends to increase over the course of each day, presumably because animals become less thirsty. To deconfound changes in relative value due to changes in value range from this effect, we alternated sessions in which we increased or decreased the range of either juice A or juice B. The number of sessions for each of the 4 possible combinations was not-predetermined with a statistical method but was comparable (ΔA→ 2ΔA, 61 sessions; ΔB → 2ΔB, 62 sessions; 2ΔA → ΔA, 49 sessions; 2ΔB → ΔB, 48 sessions). Second, within each trial block, monkeys might experience juice-specific satiety or diminishing marginal returns. Thus to isolate the behavioral effects of manipulating the value range, we ensured that in both trial blocks the animal drank the same relative amounts of the two juices. For example, if the animal drank juice A and juice B in quantity ratio 3:2 in the first block, we kept the same ratio 3:2 in the second block (see below). Third, we previously found that, all other things equal, monkeys tend to choose on any given trial the same juice they chose in the previous trial (choice hysteresis)^22^. If the relative number of trials in which the animal chooses a particular juice varies from one block to the other, choice hysteresis could introduce a systematic bias. To avoid this confound, we ensured that the relative number of choices was the same in the two trial blocks.

The relative number of choices and the relative amount drunk by the animal for each juice were controlled by adjusting the frequency with which each offer type was presented. Specifically, offers were presented pseudo-randomly in mini-blocks of 20-30 trials. To fine-tune the balance between juice A and B, we kept track of the monkey’s choices online. If the choice ratio or the relative amount of juice changed in the second block, the imbalance was corrected by adding forced choices of one of the two juices.

### Analysis of behavioral data

Monkeys’ choices generally presented a quality-quantity trade-off. If the two juices were offered in equal amounts, the animal would generally choose A (by definition). However, if sufficiently large quantities of juice B were offered against one drop of juice A, the animal would choose B. Choices were analyzed separately in each session (Exp.1) or in each trial block (Exp.2). The "choice pattern" was defined as the percentage of trials in which the animal chose juice B as a function of the log quantity ratio log(*q_B_*/*q_A_*), where *q_A_* and *q_B_* indicate the quantities of juices A and B. Each choice pattern was fitted with a sigmoid function:

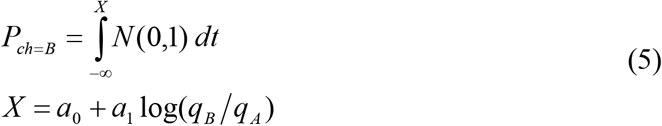

where *P_ch=B_* is the probability of choosing juice B and *N* (0,1) is the standard normal distribution. The fit was done with Matlab function glmfit and link = probit. From the fitted parameters *a_0_* and *a_1_* we defined the relative value *ρ* and the steepness of the sigmoid *η* as follows:

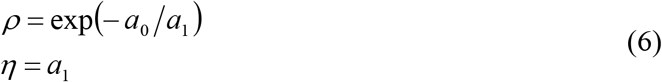

Given a set of offers, the expected payoff is directly related to *η*. In some simulations (Fig.S9), we reduced the sigmoid steepness (e.g., *a_1_* → *a_1_*/10) while keeping the relative value constant (*a_0_*/*a_1_* → *a_0_*/*a_1_*).

Notably, Exp.1 included 208 sessions. However, in some cases the choice patterns were saturated (i.e., the animal did not split decisions for any offer type, a situation referred to as "perfect separation"). In these cases, the sigmoid fit did not provide a reliable measure for *η*. Thus the analysis shown in Fig.5 and Fig.6 included only sessions for which choice patterns were not saturated (164 sessions).

In Exp.1, the minimum quantity offered for each juice was always 0, and we indicate maximum quantities with *Q_A_* and *Q_B_*. We usually set *Q_A_* and *Q_B_* to approximately satisfy *ρ Q_A_* = *Q_B_* (symmetric condition). However, this relation did not hold strictly, partly because the relative value *ρ* was determined by the animal and fluctuated from session to session. Thus to test a theoretical prediction on choice variability and value range, we computed the geometric mean value range Δ ≡ (*ρ Q_A_ Q_B_*)^1/2^ and we examined the relation between *η* and Δ. Since errors of measure affected both measures, standard regressions could not be used. We thus used Deming’s regressions^54^. Variance ratios *λ* were computed through error propagation as follows:

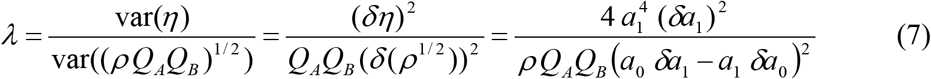

where *δρ*, *δη*, *δa*_0_ and *δa*_1_ are errors on the respective measures, and *δa*_0_ and *δa*_1_ are obtained as standard errors from the logistic regressions.

One concern was whether the relation between choice variability and value range (Fig.6) was direct or reflected some other dependency. We considered two issues. First, Fig.6 includes sessions with different juice pairs, with different typical values of *ρ*. In principle, choice variability could vary from juice pair to juice pair in a way that induces the relation observed in Fig.6. Second, for any given juice pair, sigmoid steepness (*η*), relative value (*ρ*) and value range (Δ) are all inter-related by definition (Eq.6) and because value ranges were often chosen non independently of *ρ* (in many sessions we set *Q_A_* = 3 and chose *Q_B_* such that *Q_B_* ≈ *ρ Q_A_*). Thus the relation between *η* and Δ (Fig.6) might simply reflect fluctuations in *ρ*. To address these concerns, we divided sessions in different sets based on the animal and on the juice pair. We considered only sets with ≥5 sessions, and our data included 12 such sets (6 from each monkey). We then analyzed each set of sessions separately. First, we verified that the relation between Δ and *η* held true for individual juice pairs (Fig.S5). Second, to assess whether this relation simply reflected fluctuations in *ρ*, we used multilinear regression. For each set, we regressed *η* on *ρ* and then on Δ in a stepwise way. The coefficient obtained from the second regression (*β_η_*) essentially quantified the correlation between *η* and Δ not explained by fluctuations of *ρ*.

The relation between *η* and Δ was also analyzed using alternative definitions for Δ including the simple mean Δ ≡ (*ρ Q_A_* + *Q_B_*)/2 and the log geometric mean Δ ≡ log(*ρ Q_A_ Q_B_*)/2, adjusting variance ratios accordingly. All variants of the analysis provided very similar results.

### Analysis of neuronal data

Neuronal data were collected in Exp.1. The data set included 931 cells from central OFC (area 13). The number of cells recorded was not pre-determined using statistical methods. Firing rates were analyzed in seven time windows aligned with different behavioral events (offer, go, juice). A "neuronal response" was defined as the activity of one cell in one time window as a function of the trial type. Neuronal responses were computed by averaging firing rates across trials for each trial type.

In a previous study^21^, we conducted a series of analyses to identify the variables encoded in this area. First, we submitted each neuronal response to a 3-way ANOVA (factors: position x movement direction x offer type), and we imposed a significance threshold p<0.001. In total, 505 (54%) neurons passed the criterion for factor offer type in at least one time window. Pooling neurons and time windows, 1,379 neuronal responses passed the ANOVA criterion, and subsequent analyses were restricted to this data set. Second, we defined a large number of variables including variables related to individual juices (*offer value A*, *offer value B*, *chosen value A*, *chosen value B*), other value-related variables (*chosen value*, *other value*, *value difference*, *value ratio*, *total value*), number-related variables (*chosen number*, *other number*, etc.) and choice-related variables (*chosen juice*). We performed a linear regression of each response onto each variable. If the regression slope differed significantly from zero (p<0.05), the variable was said to "explain" the response. Because variables were often correlated with each other, the same neuronal response was often explained by more than one variable. Thus for each response we also identified the variable that provided the best fit (i.e., the highest R^2^). Third, we proceeded with a variable selection analysis to identify a small subset of variables that best explained the neuronal population. We adapted two methods originally developed for multi-linear regressions in the presence of multi-collinearity, namely stepwise and best-subset^55,56^. In the stepwise method, we identified the variable and time window that provided the highest number of best fits, and removed from the data set all the responses explained by that variable. We then repeated the procedure until when the number of responses explained by additional variables was <5%. While intuitive, the stepwise procedure did not guarantee optimality. In contrast, the best-subset procedure (an exhaustive procedure) guaranteed optimality. In this case, for n = 1, 2, 3, … we computed the number of responses and the total R^2^ explained by each subset of n variables. The best subset was identified as that which explained the highest number of responses or the maximum total R^2^. In the original study, the stepwise and best-subset procedures identified the same 4 variables, namely *offer value A*, *offer value B*, *chosen value* and *chosen juice*. Fourth, we conducted a post-hoc analysis. While the explanatory power of variables included in the best subset was (by definition) higher than that of any other subset of variables, the procedure did not guarantee that this inequality was statistically significant. The post-hoc analysis addressed this issue by comparing the marginal explanatory power of each variable in the best subset with that of other, non-selected variables (binomial test). In the original study and subsequent work^34,57,58^, we found that the explanatory power of *offer value A*, *offer value B*, *chosen value* and *chosen juice* was statistically higher than that of any other variable.

The results of these analyses provided a classification for neuronal responses. Specifically, each neuronal response was assigned to the variable that explained the response and provided the highest R^2^. Thus we identified 447 *offer value*, 370 *chosen value* and 268 *chosen juice* responses. Subsequent analyses of the same data set demonstrated range adaptation^23^ and quantified noise correlations^32^.

Previous work suggested that the encoding of value in OFC was close to linear^21,23^. In this study, we conducted more detailed analyses to quantify how neuronal responses departed from linearity. Furthermore, we compared the curvature measured in neuronal responses with that measured for the cumulative distribution of the encoded values. The cumulative distribution of encoded values was taken as an important benchmark by analogy to sensory systems^2,7^. For each *offer value* response, we identified the value levels present in the session (i.e., the unique values). We then calculated the corresponding number of trials, divided it by the total number of trials in the session, and computed the corresponding cumulative distribution function (ntrials_CDF_). For each value level, we also averaged the firing rates obtained across trials. The range of offered values and the range of firing rates varied considerably from session to session and across the population (Fig.S3). Thus to compare the results obtained for different responses, we normalized offer value levels, neuronal firing rates, and ntrials_CDF_. Value levels were simply divided by the maximum value present in the session. For example, normalized values for juice B were defined as *q_B_*/*Q_B_*. For neuronal firing rates, we performed the linear regression **y** = a_0_ + a_1_ **x**, where **x** are normalized value levels. Firing rates were thus normalized with the transformation fr → (fr − a_0_)/a_1_. Similarly, for ntrials_CDF_, we performed the linear regression **y** = b_0_ + b_1_ **x**, and we normalized them with the transformation ntrials_CDF_ → (ntrials_CDF_ − b_0_)/b_1_. Examples of normalized firing rates and normalized ntrials_CDF_ are illustrated in Fig.1d,g. To estimate the overall curvature, we fit each normalized response function with a 2D polynomial and compared the quadratic coefficient (β_2_) with that obtained from fitting the corresponding normalized ntrials_CDF_. To estimate the overall S shape, we fit the normalized response function with a 3D polynomial and compared the cubic coefficient (β_3_) with that obtained for the corresponding normalized ntrials_CDF_. These measures were used for the population of *offer value* responses (Fig.2ab). Separately, we repeated these analyses for the population of *chosen value* responses (Fig.S2).

Theoretical considerations indicated that optimal response functions in our experiments would have been step functions (Fig.7bc), contrary to our observations. One concern was whether empirical response functions were optimal on average across sessions, if not for any particular session. Notably, the relative value *ρ* varied from session to session, largely because we used a variety of different juice pairs. Recent work indicates that the same neurons are associated with different juices in different sessions, with remapping dictated by the preference ranking^58^. In any given session, the optimal *offer value B* response function would have been a step function with step at x = *ρ*. However, since *ρ* varied from session to session, the resulting optimal response function would have been more gradually increasing. In fact, if the distribution of *ρ*/*Q_B_* across session had been uniform in the interval [0 1], the mean optimal response for *offer value B* neurons would have been linear. An important caveat is that the rationale that would justify linear *offer value B* responses did not hold for *offer value A* responses. In any case, we examined the distribution of *ρ*/*Q_B_* (Fig.7d). For *offer value B*, the mean optimal response function was computed as the cumulative distribution function for *ρ*/*Q_B_*.

Importantly, the data sets included in Fig.2 and Fig.S3 were originally selected based on a procedures that only considered linear encoding of value^21^. To assess the functional form of neuronal responses without bias in favor of linearity, we repeated the variable selection procedures described above including in the analyses all the variables defined in this study. The variable selection analysis was still based on linear regressions. However, firing rates were regressed on several non-linear value variables (response functions). The analysis included linear response functions (*offer A*, *offer B*), cumulative distribution functions of offer values (*offer A ntrial_CDF_*, *offer B ntrial_CDF_*), variance-equalized versions of the cumulative distribution functions of offer values (*offer A ntrial_CDF_ VE*, *offer B ntrial_CDF_ VE*; see below), optimal response functions (*offer A ORF*, *offer B ORF*), mean optimal response functions across sessions (*offer A mean(ORF)*, *offer B mean(ORF)*), and optimal response functions obtained under the assumption that the joint distribution of offers is uniform (*offer A ORF_uniform_*, *offer B ORF_uniform_*; see below). In addition, the analysis included *chosen value*, the cumulative distribution function for chosen values (*chosen value ntrial_CDF_*), a variance-equalized version of the cumulative distribution function for chosen values (*chosen value ntrial_CDF_ VE*), and variables *other value*, *value difference*, *value ratio*, *total value* and *chosen juice* (20 variables total).

We restricted the variable selection analysis to responses that passed the ANOVA criterion (N = 1,379, see above) and we regressed each neuronal response on each variable. For variance-equalized variables, we first computed the square root of the firing rates and then performed the linear regressions^59,60^. Optimal response functions for uniform joint distribution of offers (ORF_uniform_) were computed numerically in Matlab. For variable selection we used the two procedures described above, and we refer to previous work for additional details^21^. The best-subset method can be based either on the number of responses or on the total R^2^ explained by each subset. In previous studies, the two metrics provided similar results. Here we found that the results obtained based on the total R^2^ were more robust, probably because the present analysis was aimed at providing a better fit for the neuronal responses as opposed to explaining more responses. The best-subset procedures and post-hoc analyses were performed on collapsed variables^21^. The variable selection analyses were conducted twice. First, we included all the variables described above except those based on ORF_uniform_. In this case, linear response functions performed significantly better than all the other variables (Table S1). Second, we added in the analysis the variables based on ORF_uniform_. In this case, the performance of ORF_uniform_ variables was better than, but statistically indistinguishable from that of linear variables. It was significantly better than that of all the other variables (Table S2). Both analyses are described in the Results. Figs.8-10 refer to the analysis that included all 20 variables.

## References

1. Barlow, H.B. Possible principles underlying the transformations of sensory messages. in Sensory Communication (ed. Rosenblith, W.A.) 217–234 (MIT Press, Cambridge, MA, 1961).

2. Laughlin, S. A simple coding procedure enhances a neuron’s information capacity. Z Naturforsch C 36, 910–912 (1981).

3. Simoncelli, E.P. & Olshausen, B.A. Natural image statistics and neural representation. Annu Rev Neurosci 24, 1193–1216 (2001).

4. Laughlin, S.B. The role of sensory adaptation in the retina. J Exp Biol 146, 39–62 (1989).

5. Smirnakis, S.M., Berry, M.J., Warland, D.K., Bialek, W. & Meister, M. Adaptation of retinal processing to image contrast and spatial scale. Nature 386, 69–73 (1997).

6. Muller, J.R., Metha, A.B., Krauskopf, J. & Lennie, P. Rapid adaptation in visual cortex to the structure of images. Science 285, 1405–1408 (1999).

7. Brenner, N., Bialek, W. & de Ruyter van Steveninck, R. Adaptive rescaling maximizes information transmission. Neuron 26, 695–702 (2000).

8. Maravall, M., Petersen, R.S., Fairhall, A.L., Arabzadeh, E. & Diamond, M.E. Shifts in coding properties and maintenance of information transmission during adaptation in barrel cortex. PLoS Biol 5, e19 (2007).

9. Robinson, B.L. & McAlpine, D. Gain control mechanisms in the auditory pathway. Curr Opin Neurobiol 19, 402–407 (2009).

10. Liu, B., Macellaio, M.V. & Osborne, L.C. Efficient sensory cortical coding optimizes pursuit eye movements. Nat Commun 7, 12759 (2016).

11. Dan, Y., Atick, J.J. & Reid, R.C. Efficient coding of natural scenes in the lateral geniculate nucleus: experimental test of a computational theory. J Neurosci 16, 3351–3362 (1996).

12. Baddeley, R., et al. Responses of neurons in primary and inferior temporal visual cortices to natural scenes. Proc Biol Sci 264, 1775–1783 (1997).

13. Fairhall, A.L., Lewen, G.D., Bialek, W. & de Ruyter Van Steveninck, R.R. Efficiency and ambiguity in an adaptive neural code. Nature 412, 787–792 (2001).

14. Schwartz, O., Hsu, A. & Dayan, P. Space and time in visual context. Nat Rev Neurosci 8, 522–535 (2007).

15. Hildebrandt, K.J., Ronacher, B., Hennig, R.M. & Benda, J. A neural mechanism for time-window separation resolves ambiguity of adaptive coding. PLoS Biol 13, e1002096 (2015).

16. Musall, S., et al. Tactile frequency discrimination is enhanced by circumventing neocortical adaptation. Nat Neurosci 17, 1567–1573 (2014).

17. Webster, M.A. Adaptation and visual coding. J Vis 11, 1–23 (2011).

18. Padoa-Schioppa, C. Neurobiology of economic choice: a good-based model. Annu Rev Neurosci 34, 333–359 (2011).

19. Rushworth, M.F., Kolling, N., Sallet, J. & Mars, R.B. Valuation and decision-making in frontal cortex: one or many serial or parallel systems? Curr Opin Neurobiol (2012).

20. Wallis, J.D. Cross-species studies of orbitofrontal cortex and value-based decision-making. Nat Neurosci 15, 13–19 (2012).

21. Padoa-Schioppa, C. & Assad, J.A. Neurons in orbitofrontal cortex encode economic value. Nature 441, 223–226 (2006).

22. Padoa-Schioppa, C. Neuronal origins of choice variability in economic decisions. Neuron 80, 1322–1336 (2013).

23. Padoa-Schioppa, C. Range-adapting representation of economic value in the orbitofrontal cortex. J Neurosci 29, 14004–14014 (2009).

24. Cox, K.M. & Kable, J.W. BOLD subjective value signals exhibit robust range adaptation. J Neurosci 34, 16533–16543 (2014).

25. Kobayashi, S., Pinto de Carvalho, O. & Schultz, W. Adaptation of reward sensitivity in orbitofrontal neurons. J Neurosci 30, 534–544 (2010).

26. Padoa-Schioppa, C. & Rustichini, A. Rational attention and adaptive coding: a puzzle and a solution. American Economic Review: Papers and Proceedings 104, 507–513 (2014).

27. Hunt, L.T., et al. Mechanisms underlying cortical activity during value-guided choice. Nat Neurosci 15, 470–U169 (2012).

28. Kable, J.W. & Glimcher, P.W. The neurobiology of decision: consensus and controversy. Neuron 63, 733–745 (2009).

29. Krajbich, I., Armel, C. & Rangel, A. Visual fixations and the computation and comparison of value in simple choice. Nat Neurosci 13, 1292–1298 (2010).

30. Louie, K., LoFaro, T., Webb, R. & Glimcher, P.W. Dynamic divisive normalization predicts time-varying value coding in decision-related circuits. J Neurosci 34, 16046–16057 (2014).

31. Rustichini, A. & Padoa-Schioppa, C. A neuro-computational model of economic decisions. J Neurophysiol 114, 1382–1398 (2015).

32. Conen, K.E. & Padoa-Schioppa, C. Neuronal variability in orbitofrontal cortex during economic decisions. J Neurophysiol 114, 1367–1381 (2015).

33. Haefner, R.M., Gerwinn, S., Macke, J.H. & Bethge, M. Inferring decoding strategies from choice probabilities in the presence of correlated variability. Nat Neurosci 16, 235–242 (2013).

34. Padoa-Schioppa, C. & Assad, J.A. The representation of economic value in the orbitofrontal cortex is invariant for changes of menu. Nat Neurosci 11, 95–102 (2008).

35. Grace, R.C. Violations of transitivity: Implications for a theory of contextual choice. J Exp Anal Behav 60, 185–201 (1993).

36. Tversky, A. & Simonson, I. Context-dependent preferences. Management Sciences 39, 117–185 (1993).

37. Bermudez, M.A. & Schultz, W. Reward magnitude coding in primate amygdala neurons. J Neurophysiol 104, 3424–3432 (2010).

38. Cai, X. & Padoa-Schioppa, C. Neuronal encoding of subjective value in dorsal and ventral anterior cingulate cortex. J Neurosci 32, 3791–3808 (2012).

39. Tobler, P.N., Fiorillo, C.D. & Schultz, W. Adaptive coding of reward value by dopamine neurons. Science 307, 1642–1645 (2005).

40. Diederen, K.M., Spencer, T., Vestergaard, M.D., Fletcher, P.C. & Schultz, W. Adaptive prediction error coding in the human midbrain and striatum facilitates behavioral adaptation and learning efficiency. Neuron 90, 1127–1138 (2016).

41. Diederen, K.M.J. & Schultz, W. Scaling prediction errors to reward variability benefits error-driven learning in humans. Journal of Neurophysiology 114, 1628–1640 (2015).

42. Tversky, A. & Kahneman, D. The framing of decisions and the psychology of choice. Science 211, 453–458 (1981).

43. Savage, L.J. The foundations of statistics, (Dover Publications, New York, 1972).

44. Ariely, D., Loewenstein, G. & Prelec, D. ’Coherent arbitrariness’: stable demand curves without stable preferences. Q J Econ 118, 73–105 (2003).

45. Camille, N., Griffiths, C.A., Vo, K., Fellows, L.K. & Kable, J.W. Ventromedial frontal lobe damage disrupts value maximization in humans. J Neurosci 31, 7527–7532 (2011).

46. Gallagher, M., McMahan, R.W. & Schoenbaum, G. Orbitofrontal cortex and representation of incentive value in associative learning. J Neurosci 19, 6610–6614 (1999).

47. Rudebeck, P.H. & Murray, E.A. Dissociable effects of subtotal lesions within the macaque orbital prefrontal cortex on reward-guided behavior. J Neurosci 31, 10569–10578 (2011).

48. Cisek, P. Making decisions through a distributed consensus. Curr Opin Neurobiol (2012).

49. Friedrich, J. & Lengyel, M. Goal-directed decision making with spiking neurons. J Neurosci 36, 1529–1546 (2016).

50. Song, H.F., Yang, G.R. & Wang, X.J. Reward-based training of recurrent neural networks for cognitive and value-based tasks. Elife 6(2017).

51. Zhang, Z., Cheng, Z., Lin, Z., Nie, C. & Yang, T. A neural network framework for the orbitofrontal cortex and model-based reinforcement learning. bioRxiv (2017).

52. Asaad, W.F. & Eskandar, E.N. A flexible software tool for temporally-precise behavioral control in Matlab. J Neurosci Methods 174, 245–258 (2008).

53. Asaad, W.F. & Eskandar, E.N. Achieving behavioral control with millisecond resolution in a high-level programming environment. J Neurosci Methods 173, 235–240 (2008).

54. Glaister, P. Least squares revisited. Math Gaz 85, 104–107 (2001).

55. Dunn, O.J. & Clark, V. Applied statistics: analysis of variance and regression, (Wiley, New York, 1987).

56. Glantz, S.A. & Slinker, B.K. Primer of applied regression & analysis of variance, (McGraw-Hill, Medical Pub. Division, New York, 2001).

57. Cai, X. & Padoa-Schioppa, C. Contributions of orbitofrontal and lateral prefrontal cortices to economic choice and the good-to-action transformation. Neuron 81, 1140–1151 (2014).

58. Xie, J. & Padoa-Schioppa, C. Neuronal remapping and circuit persistence in economic decisions. Nat Neurosci 19, 855–861 (2016).

59. Brunel, N. & Nadal, J.P. Mutual information, Fisher information, and population coding. Neural Comput 10, 1731–1757 (1998).

60. Abbott, L.F. & Dayan, P. The effect of correlated variability on the accuracy of a population code. Neural Comput 11, 91–101 (1999).

